# SARS-CoV-2 infection of human pluripotent stem cell-derived vascular cells reveals smooth muscle cells as key mediators of vascular pathology during infection

**DOI:** 10.1101/2023.08.06.552160

**Authors:** Alexsia Richards, Andrew Khalil, Max Friesen, Troy W. Whitfield, Xinlei Gao, Tenzin Lungjangwa, Roger Kamm, Zhengpeng Wan, Lee Gehrke, David Mooney, Rudolf Jaenisch

## Abstract

Although respiratory symptoms are the most prevalent disease manifestation of infection by Severe Acute Respiratory Syndrome Coronavirus 2 (SARS-CoV-2), nearly 20% of hospitalized patients are at risk for thromboembolic events. This prothrombotic state is considered a key factor in the increased risk of stroke, which is observed clinically during both acute infection and long after symptoms clear. Here we develop a model of SARS-CoV-2 infection using human-induced pluripotent stem cell-derived endothelial cells (ECs), pericytes (PCs), and smooth muscle cells (SMCs) to recapitulate the vascular pathology associated with SARS-CoV-2 exposure. Our results demonstrate that perivascular cells, particularly SMCs, are a susceptible vascular target for SARS-CoV-2 infection. Utilizing RNA sequencing, we characterize the transcriptomic changes accompanying SARS-CoV-2 infection of SMCs, PCs, and ECs. We observe that infected SMCs shift to a pro-inflammatory state and increase the expression of key mediators of the coagulation cascade. Further, we show human ECs exposed to the secretome of infected SMCs produce hemostatic factors that contribute to vascular dysfunction, despite not being susceptible to direct infection. The findings here recapitulate observations from patient sera in human COVID-19 patients and provide mechanistic insight into the unique vascular implications of SARS-CoV-2 infection at a cellular level.

## Introduction

Over the past 20 years, human coronaviruses have been responsible for severe outbreaks worldwide, including the Severe Acute Respiratory Syndrome (SARS) outbreak of 2003 and the Middle East Respiratory Syndrome (MERS) outbreak of 2012^1^. Most recently, the global pandemic caused by the Severe Acute Respiratory Syndrome Coronavirus 2 (SARS-CoV-2) has led to over 700 million documented cases and nearly 7 million deaths as of January 2023^2^. Although the primary symptoms of SARS-CoV-2 infection are associated with the respiratory system, severe illness is also associated with devastating vascular complications^3^. The most notable of these complications is a shift to a prothrombotic state, with 20% of hospitalized patients reporting thromboembolic events and an increased risk for blood clot-related issues, such as heart attack and stroke, which can last up to a year after infection^4–7^. The frequency of these complications during and after SARS-CoV-2 infection is significantly higher than after infection with other respiratory viruses, suggesting this pathology is induced by effectors unique to SARS-CoV-2^8^.

Indicators of endothelial cell (EC) activation and dysfunction, such as oxidative stress, hyperpermeability, endothelial-to-mesenchymal transition (EndoMT), hypercoagulability, and thrombosis have all been observed in SARS-CoV-2 infected patients ^3^. However, *in vivo* direct infection of ECs remains controversial^9, 10^. Infection has, however, been detected in epithelial cells in the lower respiratory tract as well as vascular smooth muscle cells and pericytes^11–13^. Previous studies have shown that the infection of respiratory epithelial cells triggers a cascade of inflammatory responses, involving macrophages and monocytes, which can lead to widespread endothelial damage and dysfunction^8, 14^. Additionally, these immune cells can secrete a variety of pro-inflammatory cytokines and chemokines that exacerbate endothelial cell activation and dysfunction^15, 16^. It is likely the infection or activation of multiple cell types in close proximity contributes to endothelial dysfunction.

Blood clot initiation begins with endothelial cells through both extrinsic and intrinsic pathways. The extrinsic pathway is initiated through tissue factor expression in perivascular cells, including smooth muscle cells. In addition, recently published data demonstrate that vascular smooth muscle cells are crucial mediators of endothelial cell function ^17, 18^. Here, the extensive crosstalk between vascular endothelial cells and adjacent SMCs prompted us to examine if infection of SMCs cells could be the trigger for the endothelial cell dysfunction and thromboembolic events associated with infection.

In this report, we built off previously published approaches^19, 20^ to generate a new protocol to produce uniform populations of ECs and perivascular cells, including SMCs and pericytes (PCs) from human pluripotent stem cells (hPSCs). We specifically developed this approach using a serum-free and common defined media for all three cell types to isolate the effects of culture conditions away from the cellular responses to SARS-CoV-2. Using these cells, we modeled SARS-CoV-2 vascular infection under defined conditions and investigated the cell-type specific response of vascular cells to SARS-CoV-2 exposure. Our results show a preferential tropism for perivascular cells over ECs and that SMCs are likely potent sites of SARS-CoV-2 infection. In addition, exposure to SARS-CoV-2 transforms SMCs into activators of coagulation signaling in ECs which may, in turn, contribute to the devastating vascular complications observed with SARS-CoV-2 infection in human patients. Finally, our system provides a physiologically relevant platform for further studies on drug development and modeling vascular pathogenesis in a defined system.

## Results

### A fully defined, efficient, and serum-free protocol allows for differentiation, culture, and maintenance of ECs, PCs, and SMCs from hPSCs with a single common medium

We established a step-wise protocol to generate and maintain both perivascular cells (PC and SMC) and ECs in high purity (>97%) using a defined media (Fig. 1A-F). Similar to previously published approaches^19, 20^, the approach to generate ECs begins with mesendoderm specification using CHIR99021-mediated WNT activation and bone morphogenetic protein 4 (BMP4), followed by endothelial specification using vascular endothelial growth factor (VEGF), forskolin, and inhibition of platelet-derived growth factor receptor (PDGFR) and transforming growth factor beta (TGFβ) signaling, and a last stage of purification with n-cadherin inhibition, and continued PDGFR and TGFβ inhibition, and maturation with sphingosine-1-phosphate (S1P) (Fig. 1A, Supplemental Fig. 1A-B). Alternatively, after mesendoderm and early vascular specification, platelet-derived growth factor (PDGF-BB) addition with TGFβ inhibition or TGFβ addition with PDGFR inhibition allowed for the derivation of PCs and SMCs (Fig. 1B-E), respectively, in the same culture medium comprised of vascular base media comprising of Gibco Vascular SFM™, heparin, B27, epithelial growth factor (EGF), basic fibroblast growth factor (bFGF), and L-ascorbic acid-2-phosphate. All cells could be maintained and freeze-thaw recovered in the same common medium with the addition of PDGF-BB for PCs and VEGF for ECs.

**Figure 1:**
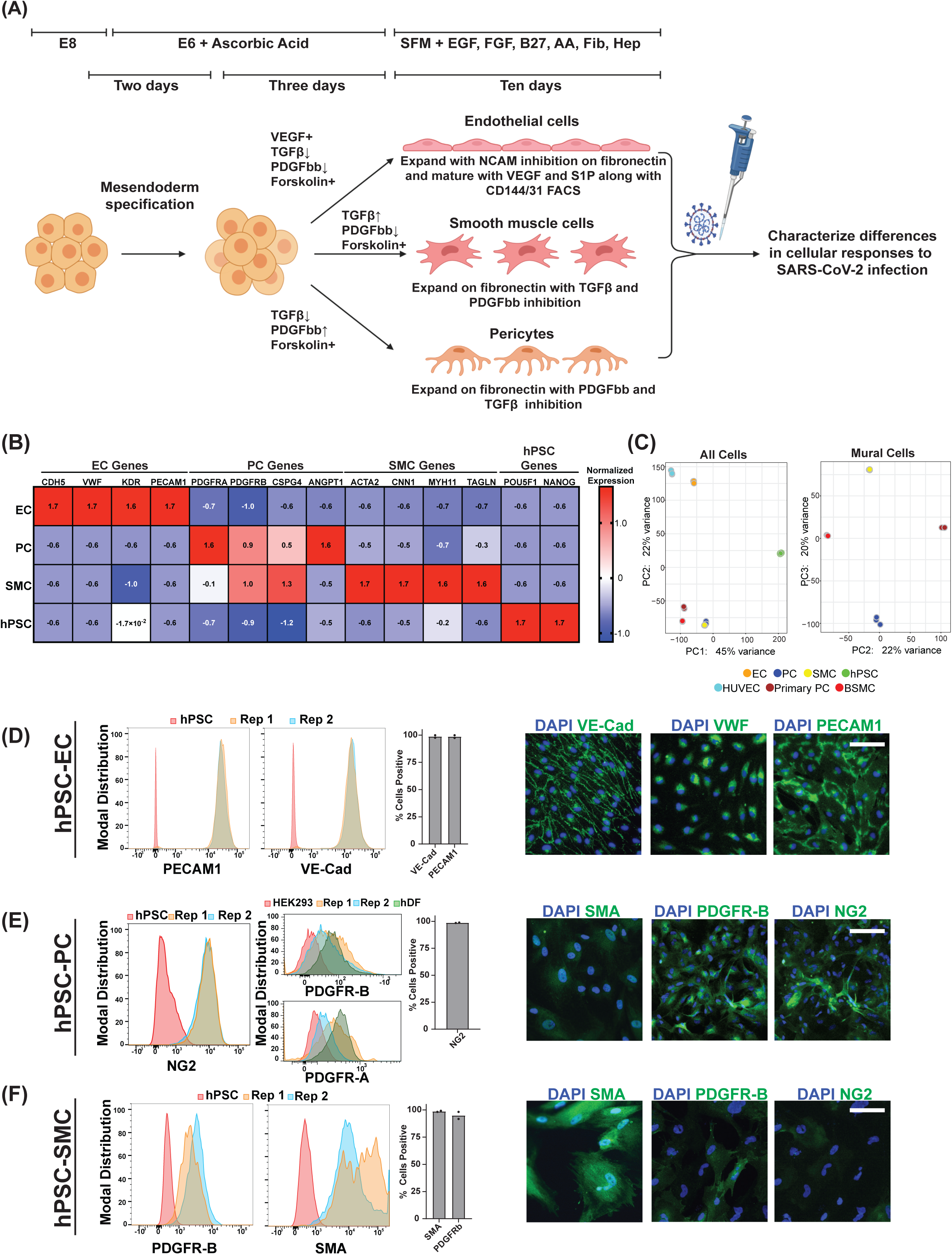
Derivation of hPSC derived vascular endothelial cells, smooth muscle cells, and pericytes. (A) Schematic of directed differentiation of hPSCs to vascular endothelial cells (ECs), smooth muscle cells (SMCs) and pericytes (PCs) (Created using BioRender). (B) Bulk RNA sequencing was performed on hPSC-derived ECs, PCs and SMCs. The normalized expression values of known EC, PC, or SMC marker genes was quantitated over three biological replicates. The values plotted represent scaled normalized expression values for each gene across samples (C) (left) Principal component analysis (PCA) on bulk-RNA sequencing data from hPSC-EC, hPSC-PCs, and hPSC-SMCs and primary endothelial cells (HUVECs), primary pericytes, and primary bronchial smooth muscle cells (BSMCs). (right) Mural cell only PCA analysis (Primary pericytes, BSMCs, hPSC-PCs, and hPSC-SMCs) (D) Expression of PECAM1, and VE-Cadherin in was quantitated by flow cytometry on hPSCs or on two independent differentiations of ECs. Data points on bar graph represent values from two independent differentiations. Expression of VE-Cadherin, VWF, and PECAM1 in hPSC-derived ECs was observed by immunofluorescence. (E) Expression of NG2 was quantitated by flow cytometry on hPSCs or on two independent differentiations of PCs. Data points on bar graph represent values from two independent differentiations. Expression of PDGFR-B, and PDGFR-A was quantitated by flowcytometry on HEK293 cells, human dermal fibroblast (hDF) and two independent PC differentiation. Expression of SMA, PDGFR-B, and NG2 in hPSC derived PCs was observed by immunofluorescence. (F) Expression of PDGFR-B or SMA was quantitated by flow cytometry on hPSCs or on two independent differentiations of SMCs. Data points on bar graph represent values from two independent differentiations. Expression of SMA, PDGFR-B, and NG2 in hPSC derived SMCs was observed by immunofluorescence. Scale bar = 50 µm for all immunofluorescence images.

For ECs, these steps resulted in >85% purity as measured by VE-Cadherin (VE-Cad/CD144) and PECAM1 (CD31) (Fig. 1D, Supplemental Fig. 1C), which could be enriched further via FACS and maintained after recovery to greater than 98% purity (Fig. 1D, Supplemental Fig. 1C). In addition, the ECs expressed canonical endothelial-related genes at levels similar to primary endothelial cells (Fig. 1B, Supplemental Fig. 1I). Our hPSC-ECs also expressed elevated levels of embryonic-restricted ETS variant transcription factor 2 (ETV2), a critical transcriptional factor expressed by endothelial cells specifically during vasculogenesis before capillary networks are formed^21^, relative to HUVECs or the starting hPSC population (Supplemental Fig. 1C). SMCs and PCs are two perivascular cell types that have been previously classified as contractile and non-contractile perivascular cells and characterized as both being PDGR positive. The delineation between the two is made through high or low levels of expression of smooth muscle actin (SMA)^19, 22, 23^ (Fig. 1D-E). Similarly, we captured these differences via gene expression in the two perivascular cell populations derived here with higher levels of contractile-related genes ACTA2 and CNN1 in SMCs and higher levels of PDGFRa and angiopoietin-1 (ANGPT1) in PCs (Fig. 1B). PC expression of PDGFRb, PDGFRa as well as the expression of the vascular adhesion molecules VCAM1 and ICAM1 was similar to levels observed in primary PCs (Supplemental Fig.1I). To further characterize our hPSC-derived vascular cell populations we performed principal component analysis (PCA) comparisons of these cell populations to each other and the primary cell counterparts (Fig. 1C). These results show that the stem cell-derived ECs group with HUVECs, while the stem cell-derived SMCs and pericytes cluster with bronchial SMCs and primary pericytes. We then further resolved the mural cell populations in separate PCA analyses without hPSCs and ECs (Fig. 1C). Euclidean distance comparisons showed that hPSC-SMCs are closer to BSMCs and hPSC-PCs to primary brain vascular pericytes (Supplemental Fig. 1G-H).

Additionally, immunohistochemistry of PCs and SMCs showed lower levels of expression in SMA and higher levels of NG2 and PDGFRb in PCs relative to SMCs (Fig. 1D-E). Importantly, relative to previously published protocols^19, 20^, both of these perivascular cells were derivable in high purity (>97%), as shown in Fig. 1D-E and Supplemental Figs. 1E-F. PCs additionally expressed intermediate levels of both PDGFRa and PDGFRb relative to fibroblasts and the starting hPSC population Fig. 1E and Supplemental Figs. 1E-F. We also examined the ability of our hPSC-derived PCs and SMCs to support microvascular network (MVN) formation with hPSC-derived ECs. Our results show that both cell populations closely associated with hPSC-EC vascular networks (Supplemental Fig. 1J). Taken together, the Euclidean distance measurements with PDGFRα/β expression comparison of the hPSC-derived SMCs and PCs to their primary cell counterparts, the examination of perivascular-specific genes such as ANGPT1, and their ability to support in vitro vascular formation collectively help to establish the authentic identity of the hPSC-derived mural cells (Fig. 1C-E & Fig. 1G-J). Furthermore, this parallel derivation in the same serum-free and defined base media specifically allowed for the comparative infection studies in one common infection medium without the confounding variables of different or undefined media formulations.

### SARS-Cov-2 demonstrates preferential tropism for human perivascular cells

To assess the potential of hPSC-derived SMCs, PCs, and ECs to be infected by SARS-CoV-2, we exposed the differentiated cells to live SARS-CoV-2 and collected media at 24 and 48 hours post-exposure. We quantified the amount of infectious virus via plaque assays and found increased infectious virus in the media from SMCs and PCs, indicating that both cell types were productively infected with SARS-COV-2 (Fig. 2A). Notably, SMCs showed the highest amount of released virus over a 48-hour infection (Fig. 2A). Conversely, exposure of ECs to SARS-CoV-2 resulted in no subsequent release of infectious virus in the media, suggesting that these cells are not productively infected with SARS-CoV-2 (Fig. 2A). Double-stranded RNA (dsRNA) is produced during SARS-CoV-2 genome replication^24^. In agreement with the plaque assay data, we detected dsRNA in a subset of hPSC-derived PCs and SMCs, indicating active viral replication, but not in ECs (Fig. 2B). To further confirm that ECs were not productively infected by SARS-CoV-2, we quantitated both positive- and negative-sense viral genome copies in ECs following exposure to SARS-CoV-2. The negative-sense genome copy is generated specifically during viral replication and thus should not be present in the input virus^24^. Our data showed that in ECs exposed to live SARS-CoV-2, levels of both positive- and negative-sense viral genomes were comparable to those in ECs exposed to heat-inactivated virus (Supplemental Fig. 3A), strongly suggesting an absence of active infection. In contrast, smooth muscle cells (SMCs) exhibited robust replication, with significantly increased levels of both positive- and negative-sense copies only after exposure to live SARS-CoV-2, and not to heat-inactivated virus (Supplemental Fig. 3A). Collectively, these data support published reports that primary endothelial cells are likely not primary targets of SARS-CoV-2 infection^9^. To determine if the expression of known SARS-CoV-2 receptors could account for the observed differences in tropism between hPSC-derived SMCs, PC, and ECs we examined the expression of ACE2, TMPRSS2, and NRP1 which have all been reported to contribute to SARS-CoV-2 entry^25, 26^. Expression of ACE2 and TMRPSS2 was higher in SMCs, and PCs, relative to ECs (Fig. 2C). ECs showed the highest expression of NRP1 amongst the three cell types (Fig 2C). This result was perhaps not surprising given the known role of NRP1 in endothelial angiogenesis^27^. NRP1 can also bind to the SARS-CoV-2 Spike protein. Critically, previous studies have shown that the expression of NRP1 alone is not sufficient to make cells susceptible to SARS-CoV-2 infection^26^. These data suggest that the expression of ACE2 and TMPRSS2 may contribute to the increased susceptibility of SMCs and PCs to SARS-CoV-2 infection. The striking differences in the susceptibility of SMCs, PCs and ECs to SARS-CoV-2 infection prompted us to compare the transcriptional response of these cell types to SARS-CoV-2 exposure.

**Figure 2:**
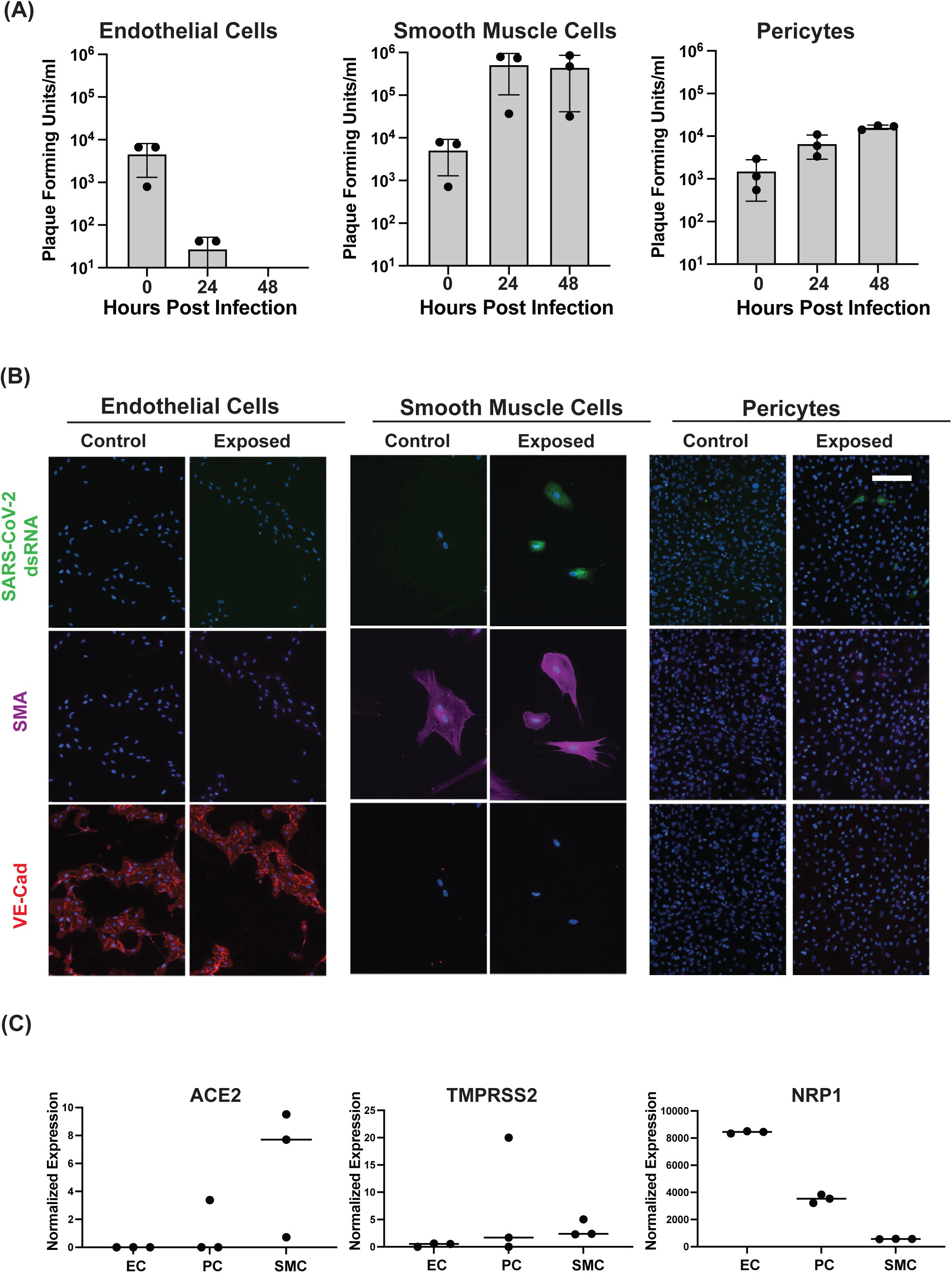
hPSC derived ECs, PCs, and SMCs show selective susceptibility to SARS-CoV-2 infection. (A) 10e^5^ hPSC-derived ECs, SMCs, and PCs were infected at an MOI of 0.1 (10e^4^ PFU) with SARS-CoV-2. The amount infectious virus released into the media at the indicated time post-infection was quantitated by plaque assay in three independent experiments. Titers of EC 48 h.p.i samples were below the limit of detection of 10 plaque forming units. Data are presented as mean values +/- SD. (B) hPSC derived ECs, SMCs, and PCs were infected at an MOI of 0.1 with SARS-CoV-2 and fixed at 48 hours post-infection. Fixed cells were stained with an antibody directed against dsRNA as well as antibodies directed against SMA or VE-Cadherin and imaged by immunofluorescence. The experiment was performed three times with similar results. Representative images from a single experiment are shown. (C) Expression of known SARS-CoV-2 receptors ACE2, TMPRSS2, and NRPP1 was quantitated by bulk sequencing on hPSC-derived ECs, PCs, and SMCs. Values shown are from three independent biological replicates. Scale bar = 50 µm for all immunofluorescence images.

### SARS-CoV-2 exposure drives divergent transcriptomic responses in human SMCs, PCs and ECs

To identify the transcriptional changes induced by SARS-CoV-2 infection, we performed bulk RNA sequencing on hPSC-derived ECs, PCs and SMCs exposed to SARS-CoV-2. Although our results showed that ECs are not productively infected with SARS-CoV-2, previous reports have demonstrated that exposure to SARS-CoV-2 proteins is sufficient to impair EC function^28^. In the model developed here, human ECs, PCs, and SMCs exposed to live SARS-CoV-2 showed significant changes in gene expression (Fig. 3A-C). Notably, only SMCs showed a strong induction of inflammatory signaling pathways following exposure to live SARS-CoV-2 (Fig 3B), ECs showed a strong upregulation of genes relating to reactive oxygen species production (Fig 3A). Surprisingly, PCs exposed to SARS-CoV-2 showed a unique transcriptional profile with few pathways significantly upregulated in infected PCs (Fig 3C).

**Figure 3:**
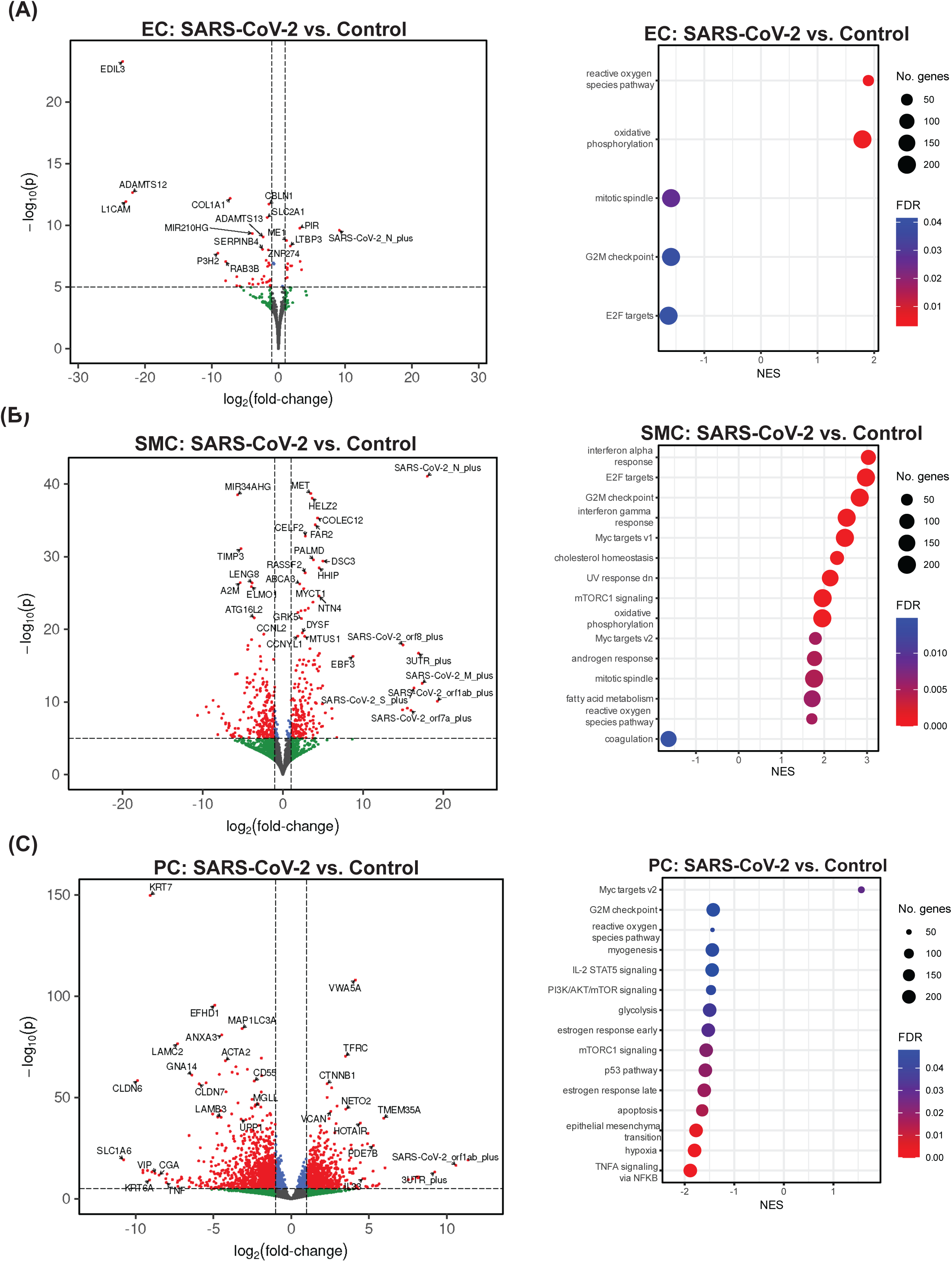
hPSC-derived ECs, SMCs and PCs display unique responses to SARS-CoV-2 exposure. Bulk RNA sequencing was performed on RNA isolated from hPSC-derived vascular ECs, PCs, and SMCs 48 hours after virus exposure (MOI =1). For all volcano plots effect sizes were estimated using DESeq2, with (two-tailed) p-values computed using the Wald statistic. A full list of differentially expressed genes can be found in the Source Data file. (A) Volcano plots showing differential gene expression in ECs exposed to live SARS-CoV-2 compared to control uninfected ECs. Gene set enrichment analysis (GSEA)^76^ was performed on differentially expressed genes to analyze the transcriptional response to infection. Dot plots show gene-sets from the Hallmark collection^34^ of the MSigDB that were enriched (FDR < 0.05). (B) Volcano plots showing differential gene expression in SMCs exposed to live SARS-CoV-2 compared to control uninfected SMCs. GSEA was performed on differentially expressed genes to analyze the transcriptional response to infection. Dot plots show genesets from the Hallmark collection^34^ of the MSigDB that were enriched (FDR < 0.05). (C) Volcano plots showing differential gene expression in PCs exposed to live SARS-CoV-2 compared to control uninfected PCs. GSEA was performed on differentially expressed genes to analyze the transcriptional response to infection. Dot plots show gene-sets from the Hallmark collection of the MSigDB that were enriched (FDR < 0.05).

The vascular complications associated with SARS-CoV-2 infection were most significant following infection with the WA-1/2020 strain of SARS-CoV-2, while later variants, which appeared in the spring of 2022 were associated with less severe clinical outcomes^29^. To determine if reduced activation of innate immune signaling in SMCs could contribute to these differences we examined the transcriptional response of SMCs to infection with the Omicron BA5.1 variant of concern (VOC). BA5.1 infection of SMCs induced inflammatory signaling, however, the enrichment of interferon-α and interferon-γ response pathways was reduced relative to the induction following infection with the WA-1/2020 strain of SARS-CoV-2 (Fig. 3A, Supplemental Fig. 2).

To identify transcriptional changes unique to productive infection with SARS-CoV-2 we compared the transcriptional response of ECs and SMCs following exposure to live SARS-CoV-2 or heat-inactivated SARS-CoV-2. Heat inactivation has previously been shown to disrupt the SARS-CoV-2 virion resulting in a non-infectious, but intact virion^30^. In ECs the changes in gene expression appeared to primarily reflect their response to SARS-CoV-2 virions, as there were few differentially expressed genes in EC exposed to live vs. heat-inactivated (HI) SARS-CoV-2 (Supplemental Fig.3A). We do observe that in ECs exposed to live SARS-CoV-2 compared to those exposed to heat-inactivated SARS-CoV-2 there is a positive enrichment of gene sets associated with metabolic processes (e.g. oxidative phosphorylation, adipogenesis, and cholesterol homeostasis), cell cycle (Myc targets), and cell death (DNA repair) pathways. (Supplemental Fig. 3A). Although we see no evidence of productive infection of ECs by SARS-CoV-2 it is possible that heat-inactivation alters the conformation of viral proteins in a way that reduces their potential to activate these pathways in ECs. Similarly, treatment with free viral spike and nucleocapsid proteins alone was also not able to recapitulate the activation of metabolic, cell cycle, or cell death pathways observed following exposure to live SARS-CoV-2 (Supplemental Fig. 3B).

SMCs responded more significantly to live SARS-CoV-2 infection with respect to differentially expressed genes, with, as expected, SARS-CoV-2 RNA being enriched in the infected SMC upregulated genes (Supplemental Fig. 4A). SMCs exposed to live SARS-CoV-2 showed a robust enrichment in the interferon-α and interferon-γ innate immune response pathways compared to SMCs exposed to heat-inactivated SARS-CoV-2 (Supplemental Fig. 4B) suggesting that productive infection amplifies inflammatory signaling. Robust inflammatory signaling continued to be observed even at 72 hours after exposure to live SARS-CoV-2 (Supplemental Fig. 4C), further supporting the hypothesis that the observed inflammatory response was due to prolonged infection and not only the result of initial activation by extracellular viral particles.

To confirm the finding that EC exposure to SARS-CoV-2 increased the generation and response to reactive oxygen species (ROS), ECs were then incubated with CellROX Green Reagent to detect the generation of ROS by immunofluorescence. A significant increase in ROS signal was detected in ECs exposed to both live and heat-inactivated SARS-CoV-2 for 48 hours (Fig. 4A-B). Induction of ROS has the potential to increase vascular permeability^31^. To determine if exposure to SARS-CoV-2 reduced endothelial cell barrier function, we measured permeability to FITC-dextran at 48 hours post-infection (Fig. 4C). 10-kDa FITC–tagged dextran was added into the apical Transwell chamber, and accumulation into the lower chamber was quantified. Our results showed that exposure to live SARS-CoV-2 resulted in nearly a 50% increase in dextran release to the lower chamber (Fig 4C). Collectively, these data both support our initial finding that SMCs are the susceptible vascular cell to SARS-CoV-2 infection as well as further show that infection and the presence of virions amplifies the divergent responses in ECs and SMCs, specifically of reactive oxygen species and induction of innate immune signaling respectively.

**Figure 4:**
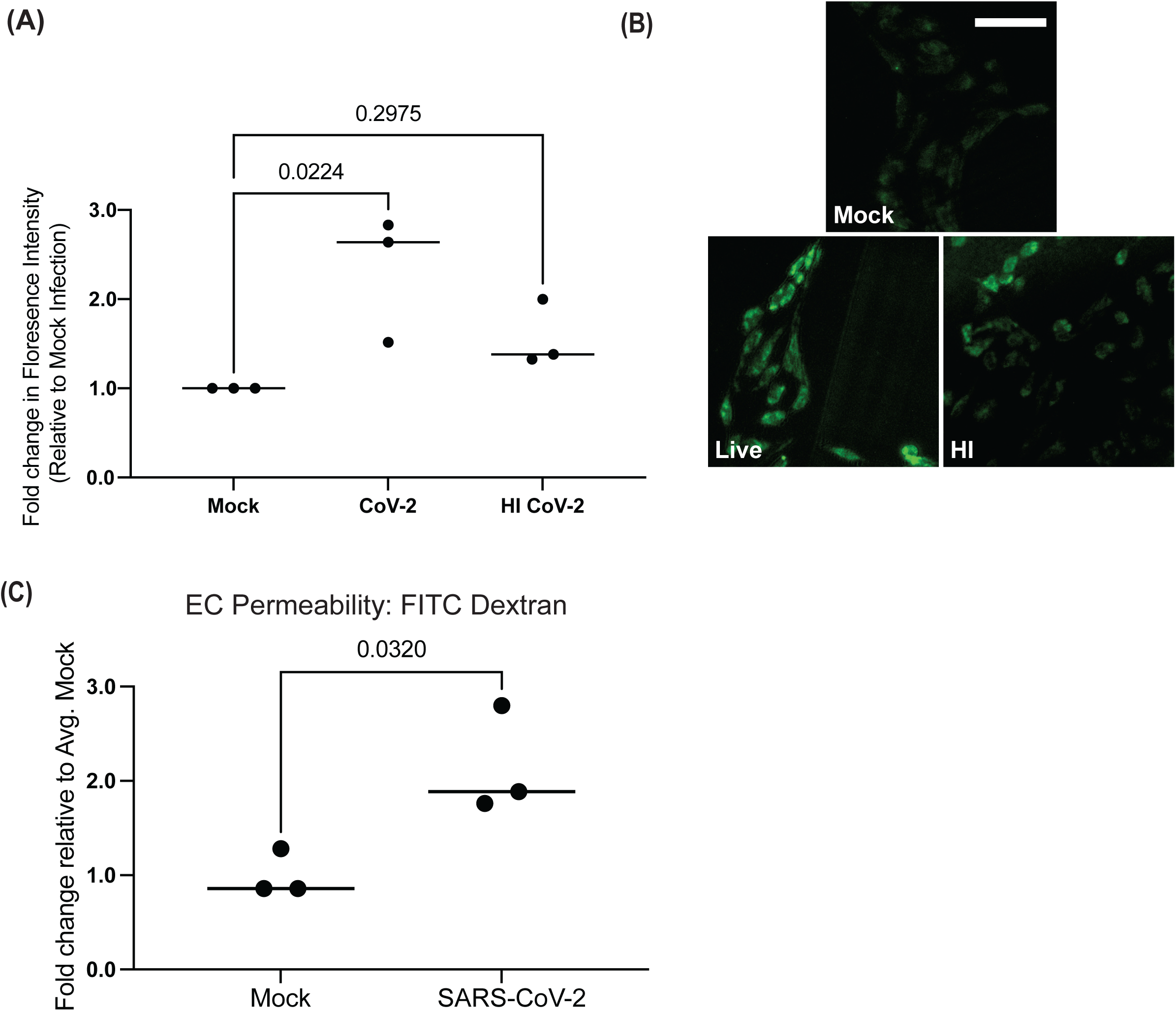
SARS-CoV-2 exposure induces reactive oxygen species production and decreases barrier function of hPSC-derived ECs. (A) ECs were mock-infected or exposed to live or heat-inactivated SARS-CoV-2 for 48 hours (MOI=1). CellROX green reagent was added at a final concentration of 5 μM and incubated for 30 minutes and cells were analyzed by fluorescence microscopy. Green fluorescent signal showed the ROS-mediated oxidation of the reagent. The fluorescence intensity was measured for five cells for each condition for three independent experiments. For each experiment the value plotted is the average fold change in florescence intensity relative to mock-infected cells for that experiment. Conditions were compared using a one-way ANOVA with Dunnett’s multiple comparisons test, with a single pooled variance. (B) Representative images of CellRox staining at 48 hours post-infection from a single experiment (Scale bar = 50 µm). (C) Transcytosis of FITC–tagged 10-kDa dextran across a monolayer hPSC-derived ECs. ECs were plated in transwells and infected with SARS-CoV-2 (MOI=1) or mock infected for 48 hours. FITC-dextran was added to the upper chamber and the fluorescence intensity of media the lower chamber was quantitated four hours after FITC-dextran addition. Three independent experiments were performed. All results are expressed as a fold change relative to the average value for mock infected ECs. Conditions were compared using an unpaired t-test.

### SMC-secreted factors induce inflammatory signaling in ECs

As shown in our data above, exposing hPSC-derived ECs directly to SARS-CoV-2 virions did not induce significant innate immune signaling (Fig. 3A, and Supplemental Fig. 3A). However, elevated inflammatory signaling is a key component of endothelial cell activation observed during SARS-CoV-2 infection^3^. We have demonstrated here that SMCs are a target for SARS-CoV-2 infection, and the crosstalk between SMCs and ECs has been previously shown to contribute to the pathology of vascular disease^17^. We hypothesized that ECs could be impacted by factors secreted by nearby infected SMCs. To test this hypothesis, we exposed ECs to media collected from infected SMCs (CoV-2 SMC CM) (Fig. 5A). In addition, we collected media from SMCs exposed to HI SARS-CoV-2 (HI SMC CM) to identify any changes in ECs that result from the SMC response to viral particles alone (Fig. 5A). Exposure of ECs to either CoV-2 SMC CM or HI SMC CM resulted in significant changes in gene expression when compared to control ECs (Fig. 5B-C, Supplemental Fig. 5).

**Figure 5:**
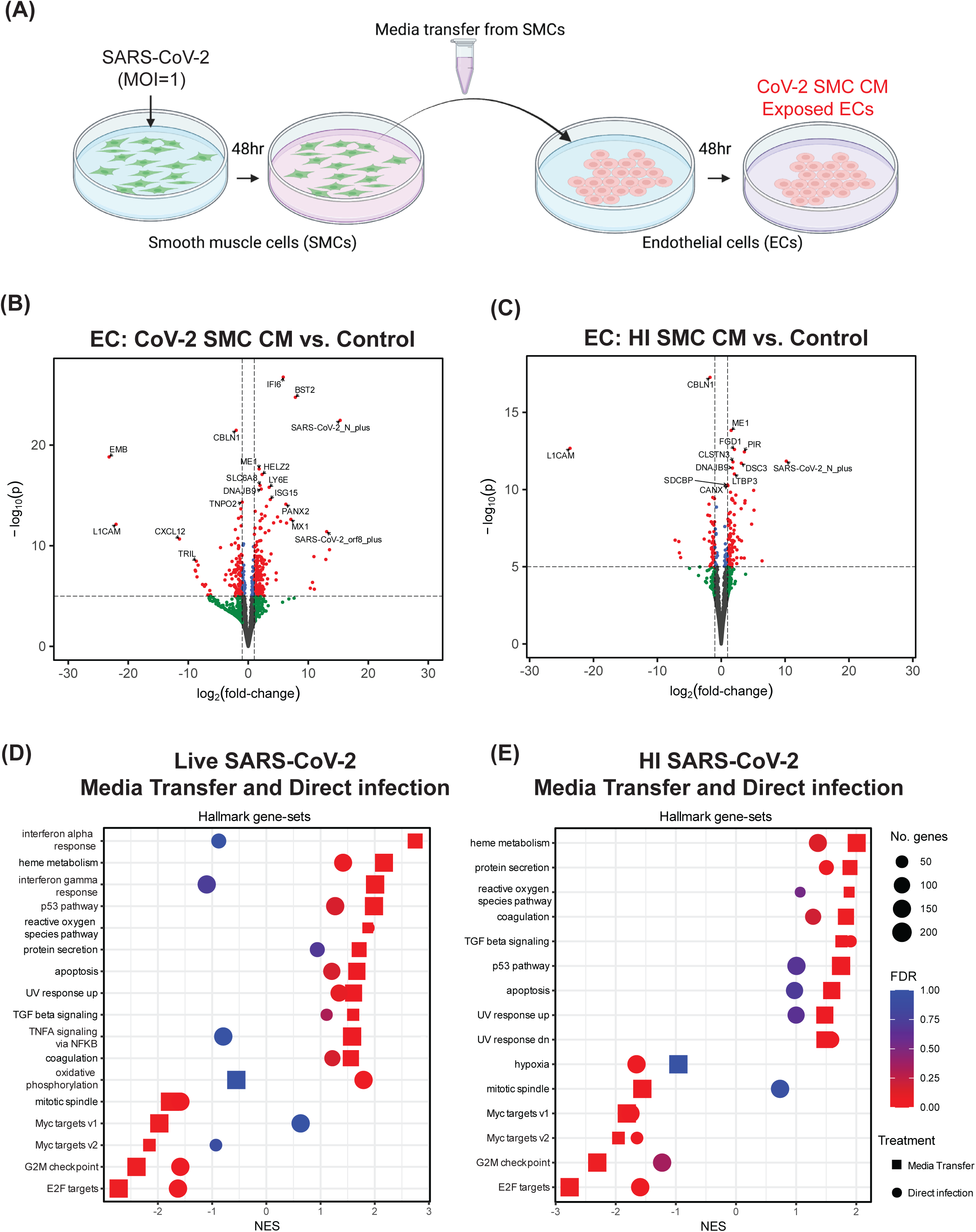
SMCs exposed to SARS-CoV-2 release factors that alter EC gene expression. **(A)** Schematic of the experiment examining the effect of SMC-secreted factors on ECs (Created using BioRender). (B-E) Bulk RNA sequencing was performed on RNA isolated from ECs 48 hours after exposure to SMC conditioned media. For all volcano plots effect sizes were estimated using DESeq2, with (two-tailed) p-values computed using the Wald statistic. A full list of differentially expressed genes can be found in the source data file. (B) Volcano plot of differentially expressed genes in ECs exposed to media from SARS-CoV-2 infected SMCs (*CoV-2 SMC CM*) compared to control ECs *(Control)*. (C) Volcano plot of differentially expressed genes in ECs treated with media from SMCs exposed to heat-inactivated SARS-CoV-2 *(HI SMC CM)* compared to control ECs *(Control)*. (D) GSEA results of ECs treated with media from SARS-CoV-2 infected SMCs compared to control ECs (*Media Transfer*) plotted together with GSEA results for ECs exposed to live SARS-CoV-2 compared to control ECs (*Direct Infection*) (E) GSEA results of ECs treated with media from SMCs exposed to heat-inactivated SARS-CoV-2 compared to control ECs (*Media Transfer*) plotted together with GSEA results for ECs exposed to live SARS-CoV-2 compared to control ECs (*Direct Infection*). In (D) and (E), the Hallmark collection^34^ of gene-sets from the MSigDB was used and gene-sets were plotted only when FDR < 0.05 for enrichment in at least one of the between-condition comparisons.

We again performed GSEA and observed that both treatment conditions demonstrated induction of pathways that could potentially contribute to vascular dysfunction via the effects on ECs by infected neighboring perivascular cells such as SMCs. As before, we observed upregulation of genes associated with reactive oxygen species (Fig. 5D-E); however, treatment with live or HI CoV-2 SMC CM increased the normalized enrichment scores for this pathway. Unlike the direct infection effects described in Fig. 3A,C and Supplemental Fig.3A, this model, which mimics the effect of SMC secreted factors on ECs, now induced inflammatory signaling pathways in ECs, including IFN-α and IFN-γ response pathways, and TNFα signaling via NF-κB (Fig. 5D-E, Supplemental Fig. 6). Notably, the induction of many genes in the interferon response pathways was significantly higher in EC exposed to CoV-2 SMC CM compared to EC exposed to HI SMC CM (Supplemental Fig. 7). Lastly, both transfer conditions resulted in further upregulation of the previously observed metabolic pathways as well as an increase in TGFβ-signaling known to regulate EndoMT^32, 33^ and apoptosis (Fig. 5D-E). Notably, exposure of ECs to SMC infection media without SMC conditioning did not result in induction of up-regulation of the inflammatory or coagulation pathways we observed with SMC conditioned media (Supplemental Fig. 8). To determine if factors secreted by SMCs infected with BA5.1 could also induce an inflammatory response in ECs we performed similar experiments using BA5.1 SMC conditioned media (BA5.1 SMC CM). While we did observe an upregulation in IFN-α and IFN-γ pathways in ECs treated with BA5.1 SMC CM, the differential expression of many of the key regulators of the innate immune response that we observed as upregulated following exposure to CoV-2 SMC CM was significantly reduced in treatment with BA5.1 SMC CM (Supplemental Figure S9, Table 1).

**Table 1:**
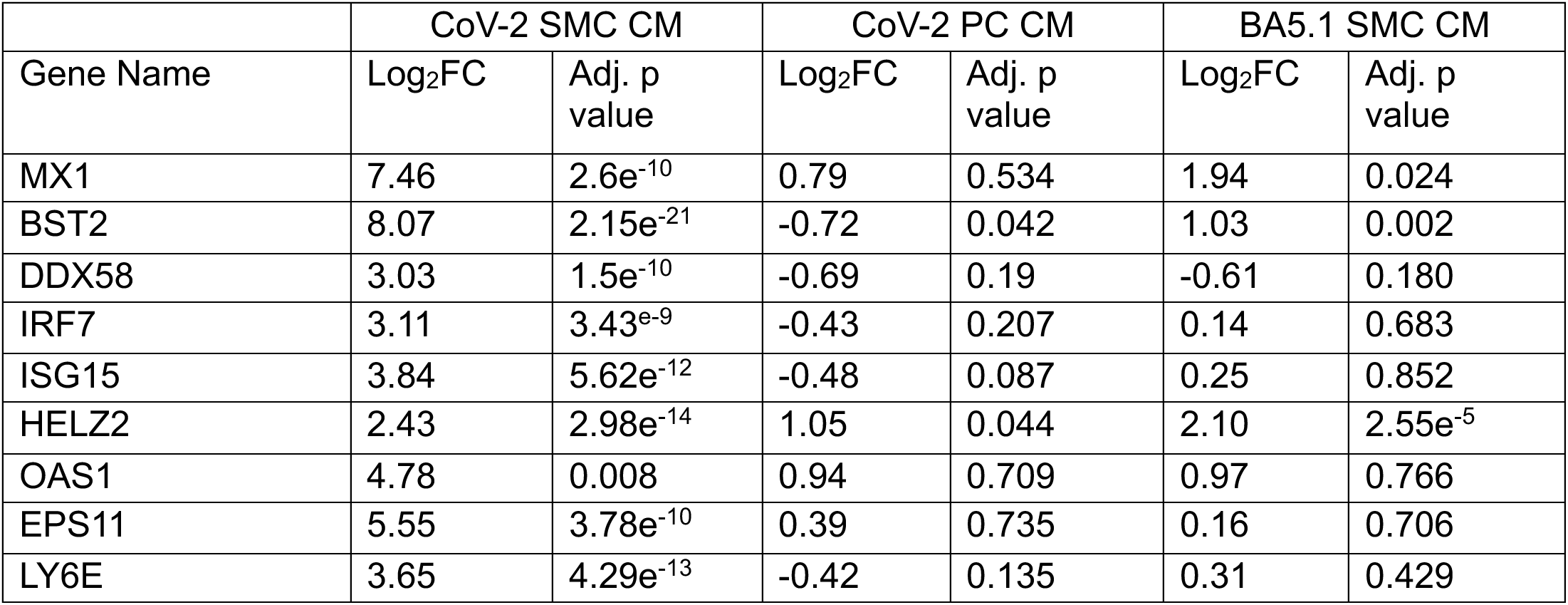
Differential expression of key regulators of innate immune signaling in ECs exposed to conditioned media from SARS-CoV-2 infected SMCs (CoV-2 SMC CM), SARS-CoV-2 infected PC (CoV-2 PC CM), or BA5.1 infected SMCs (BA5.1 SMC CM) compared to control ECs. Effect sizes were estimated using DESeq2, with (two-tailed) p-values computed using the Wald statistic. A full list of differentially expressed genes can be found in the source data file.

| Gene Name | CoV-2 SMC CM |  | CoV-2 PC CM |  | BA5.1 SMC CM |  |
| --- | --- | --- | --- | --- | --- | --- |
|  | Log<sub>2</sub>FC | Adj. p value | Log<sub>2</sub>FC | Adj. p value | Log<sub>2</sub>FC | Adj. p value |
| MX1 | 7.46 | 2.6e<sup>-10</sup> | 0.79 | 0.534 | 1.94 | 0.024 |
| BST2 | 8.07 | 2.15e<sup>-21</sup> | -0.72 | 0.042 | 1.03 | 0.002 |
| DDX58 | 3.03 | 1.5e<sup>-10</sup> | -0.69 | 0.19 | -0.61 | 0.180 |
| IRF7 | 3.11 | 3.43e<sup>-9</sup> | -0.43 | 0.207 | 0.14 | 0.683 |
| ISG15 | 3.84 | 5.62e<sup>-12</sup> | -0.48 | 0.087 | 0.25 | 0.852 |
| HELZ2 | 2.43 | 2.98e<sup>-14</sup> | 1.05 | 0.044 | 2.10 | 2.55e<sup>-5</sup> |
| OAS1 | 4.78 | 0.008 | 0.94 | 0.709 | 0.97 | 0.766 |
| EPS11 | 5.55 | 3.78e<sup>-10</sup> | 0.39 | 0.735 | 0.16 | 0.706 |
| LY6E | 3.65 | 4.29e<sup>-13</sup> | -0.42 | 0.135 | 0.31 | 0.429 |

SARS-CoV-2 productively infected pericytes; however, we did not observe induction of inflammatory signaling following infection (Fig. 3C). To determine whether factors secreted from SARS-CoV-2 infected pericytes induced inflammatory signaling in ECs we performed a parallel experiment using media conditioned by pericytes exposed to SARS-CoV-2 (CoV-2 PC CM). We observed enrichment in IFN-α and IFN-γ response pathways in CoV-2 PC CM exposed ECs compared to control ECs (Supplemental Fig. 10); however, with the exception of HELZ2 none of the major mediators of inflammatory signaling that we observed upregulated in ECs treated with CoV-2 SMC CM were significantly upregulated in ECs treated with CoV-2 PC CM (Table 1, Supplemental Fig. 6, and Supplemental Fig. 10B). Taken together, these data indicate that, while no substantial responses outside of increased reactive oxygen species-related gene signaling were detected after direct exposure, ECs can be impacted by factors released from nearby, infected perivascular cells. In particular, we observed upregulation in pathways with a potential connection to vascular dysfunction (innate immune signaling, metabolic disruption, TGFβ-signaling, and cell death) in response to SMC- and PC-secreted factors.

### SMC-secreted factors increase the expression of known coagulation pathways in ECs

In addition to the enriched pathways described above, we also observed that exposing ECs to media from infected or HI-exposed SMCs significantly upregulated the expression of multiple genes involved in coagulation (Fig. 5D-E and 6A-B). Included in the coagulation gene set^34^ was SERPINE1, a known inhibitor of plasminogen (PA-1) and a blood biomarker of which elevated plasma levels have been observed in patients with severe SARS-CoV-2 infection^35^ (Fig. 6A-B). To determine if the exposure of ECs to media from infected SMCs also increased protein secretion, we measured SERPINE1 protein levels in the media from ECs exposed to SARS-CoV-2 infected SMCs for 48 hours (CoV-2 SMC-Exposed). CoV-2 SMC-exposed ECs released significantly more SERPINE1 than ECs exposed to media from mock-infected SMCs (Mock SMC Exposed) (Fig. 6C). Similar to SERPINE1, increased plasma levels of von Willebrand Factor (vWF), a blood glycoprotein involved in clotting released from activated ECs and a common biomarker for coagulopathies^36^, has been observed in patients with severe SARS-CoV-2 infection^35, 37^. ECs exposed to media from SARS-CoV-2 infected SMCs (CoV-2 SMC Exposed) also released significantly more vWF into the media than ECs exposed to media from mock-infected SMC media (Mock SMC Exposed) (Fig. 6D). Interestingly, ECs exposed to media from BA5.1 infected SMCs did not show significant upregulation of the coagulation gene set compared to control ECs (Supplemental Fig. 9A). In addition, we observed no significant release of vWF or SERPINE1 into the media in ECs exposed to conditioned media (CM) from BA5.1 infected SMCs relative to ECs that had been exposed to CM from mock infected SMCs. (Supplemental Fig. 11). Although ECs exposed to media from SARS-CoV-2 infected PCs did show an increased expression of genes within the coagulation gene set (Supplemental Fig. 10A), we did not observe increased release of vWF or SERPINE1 into the media following exposure to CoV-2 PC CM (Supplemental Fig. 12). We propose that these data suggest that the observed coagulopathy-related vascular dysfunction observed in human patients infected with SARS-CoV-2 may be related, in part, to the paracrine effects induced by neighboring infected SMCs and the resulting increased prothrombotic signaling from ECs.

**Figure 6:**
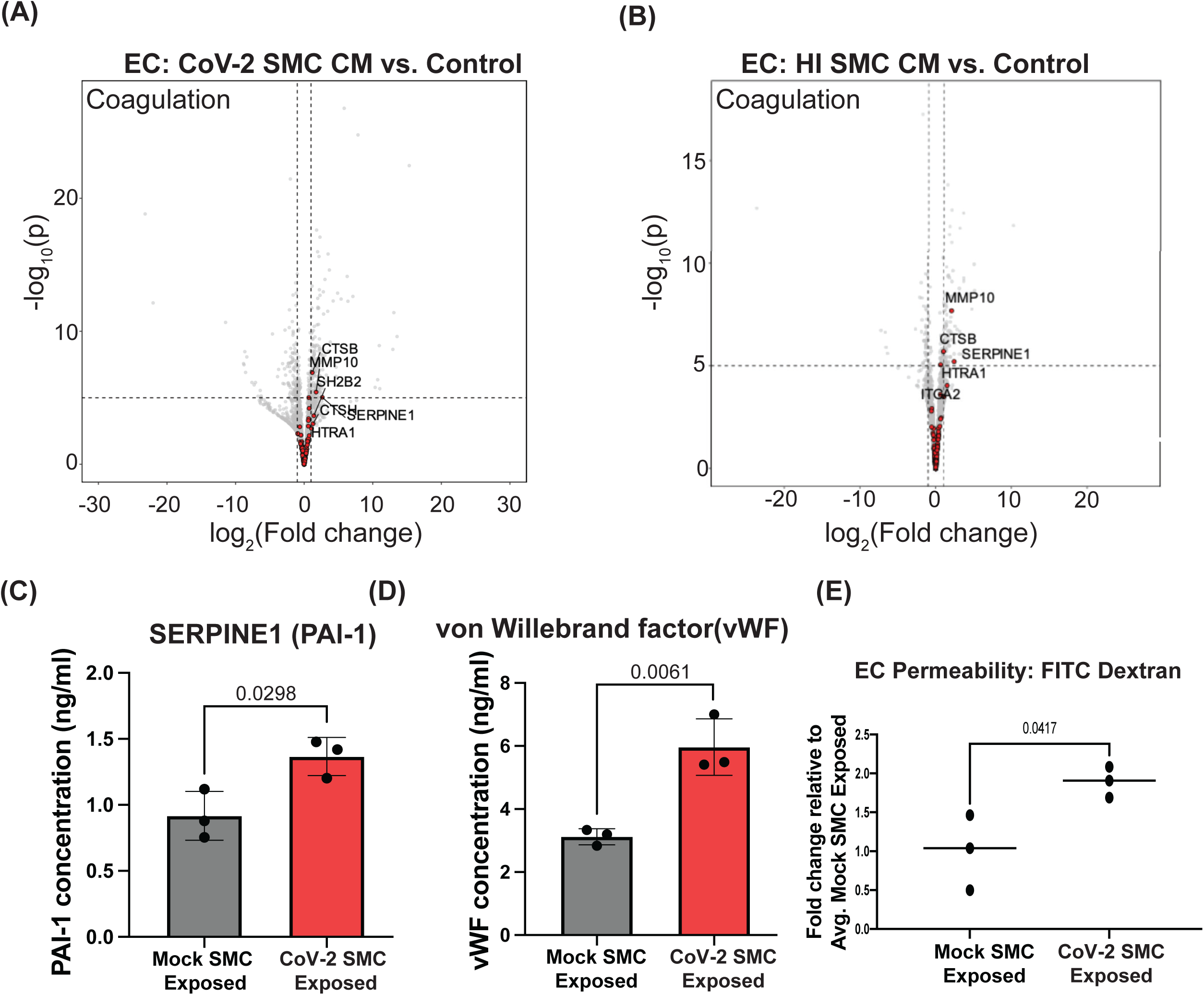
SARS-CoV-2 exposed SMCs release factors that promote clotting and reduce barrier function in ECs. (A, B) Bulk RNA sequencing was performed on RNA isolated from ECs 48 hours after exposure to SMC-conditioned media (Figure 4A) (A) Volcano plot for differential gene expression in ECs exposed to media from SARS-CoV-2 infected SMCs (*CoV-2 SMC CM*) compared to control ECs *(Control)*. For all volcano plots effect sizes were estimated using DESeq2, with (two-tailed) p-values computed using the Wald statistic. A full list of differentially expressed genes can be found in the source data file. (B) Volcano plot for differential gene expression in ECs exposed to media from SMCs exposed to heat-inactivated SARS-CoV-2 *(HI SMC CM)* compared to control ECs *(Control)*. In both (A) and (B), genes highlighted in red correspond to the “coagulation” gene-set from the Hallmark collection^34^ of the MSigDB, with GSEA “leading edge” genes labeled by name. (C-E) ECs were exposed to media from SARS-CoV-2 infected SMCs (*CoV-2 SMC Exposed*) or exposed to media from mock infected SMCs *(Mock SMC Exposed)* for 48 hours. (C) The amount of SERPINE1(PAI-1) ECs released into the media 48 hours after CM exposure was quantitated by ELISA. Media was collected from three independent experiments and analyzed in parallel. Bar graph is the average value for each condition and data points show the individual values from each experiment. Error bar shows +/- SD. Conditions were compared using an unpaired t-test. (D) The amount of von Willibrand Factor (vWF) ECs released into the media 48 hours after CM exposure was quantitated by ELISA. Media was collected from three independent experiments and analyzed in parallel. Bar graph is the average value for each condition and data points show the individual values from each experiment. Error bar shows +/- SD. Conditions were compared using an unpaired t-test. (E) Transcytosis of FITC–tagged 10-kDa dextran across a monolayer hPSC-derived ECs. ECs were plated in transwells and exposed to media from SARS-CoV-2 infected SMCs (*CoV-2 SMC CM*) or media from mock infected SMCs (*Mock SMC Exposed*) for 48 hours. FITC-dextran was added to the upper chamber and the fluorescence intensity of media the lower chamber was quantitated four hours after FITC-dextran addition. Three independent experiments were performed. All results are expressed as a fold change relative to the average value for mock-exposed ECs. Each dot represents the value from an independent experiment and lines show mean value across all three experiments. Conditions were compared using an unpaired t-test.

Infection of SMCs with SARS-CoV-2 results in significant induction of IFN-α and IFN-γ signaling pathways (Fig. 3, Supplemental Fig. 4). To determine if activation of these inflammatory pathways could contribute to the release of factors that promote activation of coagulation pathways in ECs, SMCs were treated with IFN-α or IFN-γ for 24 hours. The media was then replaced with fresh media without cytokines for 48 hours, at which point the media was collected. ECs were then incubated with IFN-α or IFN-γ SMC-conditioned media and the levels of vWF or SERPINE1 released into media were quantitated relative to ECs treated with media conditioned by untreated SMCs or PCs. ECs exposed to media from IFN-α treated SMCs showed a significant increase in the release of vWF as well as increased release of SERPINE1 (Supplemental Fig. 13A). We performed similar experiments using media conditioned by IFN-α or IFN-γ treated PCs. Treatment of ECs with media from IFN-α or IFN-γ PCs did not cause an increase in release of vWF or SERPINE1 (Supplemental Fig. 13B). Collectively, these findings suggest that activation of inflammatory signaling in SMCs may contribute to the release of factors that promote activation of coagulation cascades in ECs.

### Infected SMCs release factors that increase brain microvascular endothelial cell permeability

A recent study has shown disruption of the blood-brain barrier in patients during acute SARS-CoV-2 infection as well as in patients suffering from long COVID cognitive impairment ^38^. hPSC-derived brain microvascular endothelial cells (hBMECs) recapitulate much of the unique physiology of the brain microvascular endothelial cells in the blood-brain barrier ^39, 40^. As we observed clear dysfunction in the more peripheral-like ECs after exposure to factors secreted by infected perivascular cells, we modeled exposure of hBMECs to factors released from infected perivascular cells. Similar to the peripheral ECs, hBMECs exposed to live virus for 24 or 72 hours and stained with an antibody against dsRNA showed little to no infection, indicated by rare dsRNA-positive cells at the 72-hour time point (Supplement Fig. 14A). These data are in agreement with a previous report that showed a low level of infection by 72 hours in primary BMECs^41^. As our findings here have indicated (Fig. 5D and Supplemental Fig. 6) infected SMCs release factors that promote inflammatory signaling in ECs, we hypothesized that similar signaling would result in decreased barrier integrity in hBMECs. Transendothelial electrical resistance (TEER) provides a quantitative measurement of junctional tightness^42^. TEER was measured in hBMECs that were plated in transwell plates and exposed to media from either mock-infected SMCs (Mock CM-Exposed) or SMCs infected with live SARS-CoV-2 (CoV-2 SMC-Exposed) or heat-inactivated SARS-CoV-2 (HI SMC Exposed) over the course of 72 hours. We observed a significant reduction in TEER values over time in CoV-2 SMC Exposed hBMECs (Supplemental Fig. 14B), whereas no reduction was observed in HI SMC Exposed hBMECs (Supplemental Fig. 14C).

### hPSC-derived SMCs increase tissue factor expression and activity in response to SARS-CoV-2 infection and constitute a platform for identifying targeted antiviral therapies

Tissue factor (TF) is a key initiator of the extrinsic blood coagulation pathway expressed in vascular smooth muscle cells following injury^43^, and in the lungs of patients with severe SARS-CoV-2 infection^44,45^. To determine if SARS-CoV-2 infection of SMCs resulted in increased TF expression, SMCs were exposed to either live SARS-CoV-2 or heat-inactivated SARS-CoV-2 and stained with an antibody against TF. Exposure to live SARS-CoV-2 resulted in an increase in TF staining in SMCs (Fig. 7A-B), which appeared to correlate with dsRNA positive cells (Supplemental Fig. 15). Quantifying the average intensity of SMCs expressing TF, we observed that this increase was specific to live SARS-CoV-2 infection (Fig. 7C). To further determine whether the increased surface expression of TF indicated functional activation, we employed an enzyme-linked immunosorbent assay (ELISA) to quantitate TF activity from lysate of SMCs exposed to live or heat-inactivated SARS-CoV-2. We observed significantly higher TF activity in the lysate from SMCs infected with SARS-CoV-2 (Fig. 7D). Collectively, these data suggest that the upregulation of TF is a direct effect of infection and gives additional cellular mechanistic insight into how SARS-CoV-2 infection and the tropism shown here might cause the unique vascular dysfunction observed in COVID-19 patients.

**Figure 7:**
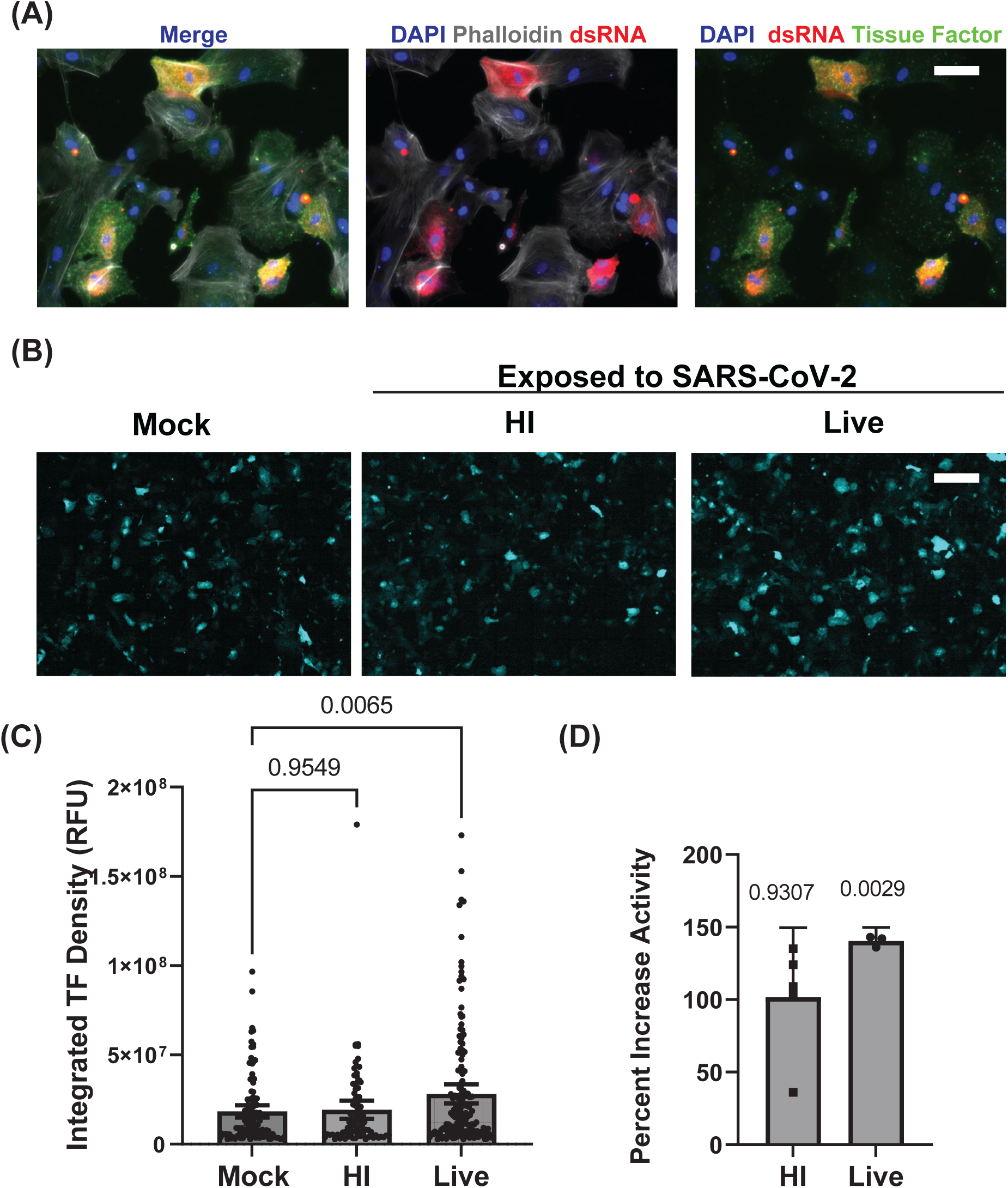
SARS-CoV-2 infection of SMCs increases tissue factor expression and activity. (A) SMCs were infected with SARS-CoV-2 for 48 hours (MOI=1) and stained with antibodies against dsRNA and tissue factor. Staining of actin with phalloidin was included to mark cell bodies. (B) Immunofluorescence staining for tissue factor in hPSC-derived SMCs exposed to either live or heat-inactivated (HI) SARS-CoV-2 or uninfected control hPSC-derived SMCs. Cells were fixed at 48 hours post-infection. Scale bar = 200 µm. The experiment was performed three times with similar results. (C) Quantitation of tissue factor staining shown in (B). Bar graph shows mean value and error bar shows 95% confidence interval. Conditions were compared using a one-way ANOVA with Dunnett’s multiple comparisons test, with a single pooled variance. (D)Tissue factor activity in cell lysate from SMCs exposed to live SARS-CoV-2 or HI SARS-CoV-2 (MOI=1) for 48 hours was quantitated by ELISA and is reported as a percent increase compared to control uninfected SMCs. Scale bar = 50 µm for all immunofluorescence images. Activity levels were analyzed for three independent experiments for live SARS-CoV-2 treatment and five independent experiments for HI SARS-CoV-2. Levels were normalized to the average value for control SMCs for that experiment. Bar graph shows mean value and error bar shows 95% confidence interval. Each condition was compared to 100% activity for control SMCs using a one sample t test.

Given that our data here suggest that infection of SMCs may be one of the major initiating factors in the development of the vascular pathology associated with infection, we sought to determine whether inhibiting known viral infection pathways in perivascular cells would blunt this as a proof-of-principle of this model’s use in drug development in infectious disease with vascular complications. The sphingolipid synthesis pathway has been shown to play a critical role in the replication of viruses and regulation of the innate immune response^46^ and is involved in the development and maturation of blood vessels^47, 48^. Modulation of this pathway has been proposed as a potential therapeutic treatment for severe infection^49^. To determine if inhibition of sphingolipid signaling in SMCs could reduce SARS-CoV-2 infection, SMCs were treated with N,N-Dimethyl-D-erythro-sphingosine (DMS), which inhibits the phosphorylation of sphingosine to form sphingosine 1-phosphate. Treatment with DMS resulted in a dose-dependent reduction of infectious virus production in SMCs, which could not be accounted for by the toxicity of the drug (Supplemental Fig. 16A). We compared the transcriptional response of SARS-CoV-2 infected SMCs with and without treatment with DMS during infection. We observed increased activation of inflammatory pathways in DMS-treated SARS-CoV-2 infected SMCs, which may in part, account for the reduced level of viral infection (Supplemental Fig. 16A-B). To determine if EC viability was impacted by treatment with DMS at a concentration that reduced viral infection in SMCs while not reducing SMC viability (Supplemental Fig. 16A), we additionally performed the CellTiter Glo viability assay on ECs 72 hours after DMS exposure and observed no reduction in EC viability relative to control DMSO exposed ECs (Supplemental Fig. 16C).

## Discussion

The SARS-CoV-2 pandemic has impacted millions of people worldwide. The vascular complications that arise from severe infection continue to devastate patients long after the peak of infection. One year after infection, patients who were infected with SARS-CoV-2 are 63% more likely to have cardiovascular issues and have a 52% increased risk of stroke^5^. In this report, we present a model system to study the tropism and pathology associated with SARS-CoV-2 vascular infection. Our data demonstrate that within the vasculature, perivascular cells, including pericytes and smooth muscle cells, are major targets for SARS-CoV-2 infection. Also consistent with previous reports^9^, our results here show no evidence of productive viral infection in human ECs. These data are particularly relevant in light of recent studies using patient biopsy samples that showed evidence of the spread of infection through the bloodstream during an early viremic phase^12^. This systemic spread likely results in virus throughout the body following infection of the respiratory tract^12^.

To enable this study, we developed a new, defined, and serum-free protocol to derive endothelial cells and multiple mural cell populations from hPSCs using a single common medium. This approach produced high-purity vascular cells that expressed canonical markers corresponding to their respective vascular identities^50^. Specifically, the perivascular cell populations, representing pericytes and smooth muscle cells, exhibited varying levels of RNA expression of key markers such as PDGFR-A, PDGFR-B, NG2, TAGLN, and SMA^50^ While specific definitions of perivascular cell identities remain important areas of investigation in vascular biology, critically, these cells were functionally capable of supporting vascular network formation via hPSC-ECs when exposed to flow (Supplemental Fig. 1J) and expressed specific makers associated with vessel support and maintenance such as ANGPT1 ^51, 52^(Fig. 1). Moreover, PCA of RNA sequencing of these cells and Euclidean distance comparisons with their primary counterparts showed that the SMCs clustered with primary bronchial smooth muscle cells and PCs with primary pericytes, respectively (Fig. 1B). The additional criteria of generating these populations of cells in a common and standardized defined medium for all cell types enabled the comparisons between these cell types under uniform infection culture conditions and, combined with the ability to form vascular networks in these defined conditions, collectively illustrate the utility of this new derivation method for the broad study of vascular biology using human pluripotent stem cells.

While we did not observe any productive infection of ECs, the RNA seq data did show perturbation of EC gene expression by exposure to SARS-CoV-2 (Fig. 3A), such as perturbations expression of in EGF-like repeats and discoidin I-like domain genes that play a significant role in endothelial cell adhesion and angiogenesis^53^. Specifically, our results showed that exposure of ECs to SARS-CoV-2 resulted in significant downregulation of EDIL3 (Fig. 3A). Consistent with this hypothesis, we observed reduced EC barrier function following exposure to SARS-CoV-2 (Fig. 4C). To better understand the direct effects on vascular endothelium from SARS-CoV-2 exposure in the absence of infection, we exposed ECs to both live and HI SARS-CoV-2 particles. The absence of significant differences following live versus HI exposure in ECs suggests the direct impact on vascular endothelium could be due to interactions between ECs and the viral proteins present on the surface of the viral particle. This finding is in line with a previous study in hamsters using pseudotyped virus expressing SARS-CoV-2 spike protein^28^. Lastly, these effects were not observed with free recombinant nucleocapsid or spike protein, indicating viral particle presentation of these proteins may be required (Supplemental Fig.3).

Our comparison of the transcriptional response in smooth muscle cells (SMCs) to live versus heat-inactivated SARS-CoV-2 revealed the specific upregulation of several inflammatory genes, including HELZ2, MX1, and IFI6, during active infection. These genes play key roles in orchestrating immune responses to viral infections and are notably upregulated in patients with severe COVID-19^54–56^. The selective elevation of these genes following exposure to live virus suggests that active SARS-CoV-2 infection enhances antiviral signaling in SMCs beyond what is triggered by viral proteins alone.

HELZ2, MX1, and IFI6 are also induced in endothelial cells (ECs) after exposure to factors secreted from SARS-CoV-2-infected SMCs. Since ECs are not directly infected by SARS-CoV-2, this data implies that inflammatory signals from neighboring SMCs are sufficient to activate expression in ECs. All three genes are interferon-stimulated genes (ISGs), whose expression is induced by interferon (IFN) signaling^57–59^, we hypothesize that infected SMCs may secrete IFNs, which then activate these antiviral pathways in nearby ECs.

Consistent with *in vivo* findings^60, 61^, our data support perivascular cells as sites of active SARS-CoV-2 infection. An important question is how viral infection could spread from the initial site of infection in the respiratory tract to perivascular cells in the absence of endothelial cell infection. During severe SARS-CoV-2 infection the virus spreads systemically through the bloodstream to peripheral sites of infection. Our data show weakened EC barrier function following exposure to SARS-CoV-2, which is consistent with patient data as well as in vitro studies ^8, 62, 63^. The specific observed reduction in EIDL3 levels in our data suggests a possible specific mechanism where EC exposure to SARS-CoV-2 particles may have the potential to impair barrier functions via weakened cell-cell adhesion and reduced angiogenesis, contributing to extravascular dissemination of the virus during COVID19 and a route to mural cell infection. We also observed the activation of ROS signaling in ECs following SARS-CoV-2 exposure, which may further contribute to reduced barrier function and increased vascular leakage. Taken together, the results here demonstrate that EC exposure to SARS-CoV-2 elicits numerous putative mechanisms of changes in gene expression and cellular function that lead to a reduction in barrier integrity and provide an access point for the virus to contact perivascular cells and disseminate into tissues, contributing to the collective understanding of COVID19 pathologies.

In identifying the sites of vascular infection, we also observed that only a subset of SMCs and PCs were positive for dsRNA staining (Supplemental Fig. 1 & Supplemental Fig. 7). This may be due to heterogeneity in receptor expression, or alternatively, the viral genomic replication that produces dsRNA may be quickly dampened by the host cell’s innate immune responses^64^. *In vivo,* it is likely that only a small percentage of interactions between viral particles and cells result in productive infection due to both the production of defective viral particles ^65^ as well as interactions of infectious viral particles with cells that are not susceptible to SARS-CoV-2 infection. However, our results comparing the transcriptional response of SMCs and ECs exposed to live virus versus heat-inactivated virus allowed us to discriminate between transcriptional changes due to productive infection and changes induced via exposure to viral particles without downstream infection. Our results suggest that the overall SMC response may be sufficient to promote vascular dysfunction, and they prompted us to expand our model to mimic the exposure of ECs to factors released from infected SMCs.

Contrary to the limited dysfunction beyond endothelial ROS responses observed in our initial model of direct infection, the new model showed significant dysregulation of vascular gene pathways in ECs. Specifically, we observed increased expression of genes in interferon response pathways, further reactive oxygen species, and significant cell death in ECs exposed to the SMC-secreted milieu. Moreover, our RNA-seq data showed that SMCs exposed to heat-inactivated SARS-CoV-2 alone, which cannot initiate infection, release factors that promote coagulation signaling in ECs, suggesting that exposure of SMCs to circulating viral particles may be sufficient for activation of coagulation pathways through a SMC dependent mechanism. Lastly, the conditioned media model provided mechanistic insights into how neighboring SARS-CoV-2 infected perivascular cells can contribute to vascular dysfunction, such as leakage and edema through decreasing barrier function.

To assess the in vivo relevance of our model system, we explored several pathways that could underlie the vascular complications seen in COVID-19, such as hypercoagulability and bleeding disorders leading to exsanguination—events our data also detected^8, 35, 37, 66, 67^. Notably, we observed an upregulation of interferon-stimulated genes and coagulation mediators like SERPINE1 (encoding for plasminogen activator inhibitor-1) and vWF in ECs exposed to conditioned media from SARS-CoV-2-infected SMCs. These are both key factors known to mediate thromboembolic events^35^ and have been observed at elevated levels in severe COVID-19 patients associated with high thrombotic risk^35, 37^ suggesting fidelity of the model findings. In addition to this EC dysfunction driven by paracrine signaling, infected SMCs in our model showed increased Tissue Factor activity, a primary driver of the intrinsic coagulation cascade^43, 45^, further aligning with in vivo reports of its upregulation in lung tissue from severe COVID-19 cases^45^ and persistent systemic inflammation^44^. Beyond SMC-driven effects, our model indicated that SARS-CoV-2 particles directly impair EC barrier function even without productive infection, potentially explaining the paradoxical bleeding and exsanguination observed in severe cases^66^. We identified significant disruptions in ROS signaling in ECs following viral exposure, consistent with increased vascular permeability seen in severe COVID-19^63^ and supported by elevated NADPH oxidase activity in the microvasculature of patients^68^. Our concurrent observation of EIDL3 downregulation, with its potential role in weakening tight junctions and compromising vascular integrity^53^, suggests a novel mechanism by which SARS-CoV-2 may induce EC dysfunction. Given the alignment of our collective findings with established observations in COVID-19 patients, further studies using human vascular tissue are warranted to validate these possible EIDL3-dependent mechanisms.

The approach detailed here describes a highly efficient manner for deriving pure perivascular and endothelial cells from hPSCs in a defined and common media for use in disease modeling. Using the defined and serum-free conditions for all cell types, this model could be used to study the tropism of SARS-CoV-2 on cells of the vasculature. The model and results described here add to the existing literature that perivascular cells, not the endothelium, are the target sites of infection. However, viral particles themselves can alter the state of ECs in the absence of infection. Additionally, the defined common medium format for all cell types allowed us to perform a set of conditioned media experiments without the complications of different media compositions and then probe the outcomes of changes in the EC microenvironment from infection of neighboring perivascular cells. This approach ultimately provided a potential intrinsic and extrinsic mechanism for coagulopathies associated with SARS-CoV-2 and paradoxical vascular leakage and edema. In summary, our findings provide a highly efficient derivation of a fully defined and serum-free hPSC vascular model and add to the knowledge of the relationship between the unique vascular complications associated with SARS-CoV-2 infection and COVID-19 patients.

## Methods

### Ethics

All research presented in this manuscript complied with all relevant ethical regulations involving the use of human pluripotent stem cells and was approved by the Committee on the Use of Humans as Experimental Subjects (COUHES) and assigned IRB protocol number #0612002068.

### hPSC lines and maintenance

H1 (WA01) embryonic stem cells were obtained from WiCell. hPSCs were maintained in feeder-free conditions in mTeSR Plus media (StemCell Technologies) on Matrigel (Corning) in 6-well tissue culture dishes. For passaging, hPSCs were detached as clumps using Versene Solution (Thermo Fisher Scientific) and replated at a ratio of 1:8-1:10.

### Directed differentiation of hPSCs to endothelial cells

All media recipes are described in Supplemental Table 2. H1 hPSCs were cultured in E8 media (Thermo Fisher Scientific) and dissociated using Accutase (Thermo Fisher Scientific) into a single cell suspension and plated at 15,000 cells/cm^2^ in E8 media with Y-27632 (10μM) and onto Matrigel-coated tissue culture plates. Y-27632 was removed after 24 hours, and cells were cultured in E8 media until they reached approximately 70% confluency, at which point the media was replaced with MelM media; this is considered day 0. On day 2, the media was replaced with EC1 media. On day 5, the media was replaced with EC2 media for 24 hours, and then cells were passaged 1:1 onto fibronectin (FC01010MG, Fisher Scientific) coated tissue culture plates in EC3 media. Plates were coated with fibronectin at 10ug/ml for 30 minutes at 37°C prior to the addition of cells. After 48 hours, media was replaced with EC4 media for 5 days and then sorted for VE-Cadherin and PECAM1 double-positive cells using FACS. Sorted cells were plated back onto fibronectin and cultured for an additional 5 days in EC4 before cryopreservation or extended expansion in EC5 medium.

### Directed differentiation of hPSCs to pericytes

All media recipes are described in Supplemental Table 2. H1 hPSCs were cultured in E8 media (Thermo Fisher Scientific) and dissociated using Accutase (Thermo Fisher Scientific) into a single cell suspension and plated at 15,000 cells/cm^2^ in E8 media with Y-27632 (10μM). Y-27632 was removed after 24 hours and cells cultured in E8 media until they reached approximately 70% confluency at which point the media was replaced with MelM media, this is considered day 0. On day 2 the media was replaced with PC/SMC1 media. On day 5 the media was replaced with PC/SMC2 media for 24 hours and then cells were passaged 1:3 on to fibronectin (FC01010MG, Fisher Scientific) coated tissue culture plates in PC3 media. Plates were coated with fibronectin at 10ug/ml for 30 minutes at 37°C prior to the addition of cells. After 48 hours media was replaced with PC4 media. After 48 hours the media was replaced with fresh PC4 and cells kept in culture for 7 days prior to freezing.

### Directed differentiation of hPSCs to smooth muscle cells

All media recipes are described in Supplemental Table 2. H1 hPSCs were cultured in E8 media (Thermo Fisher Scientific) and dissociated using Accutase (Thermo Fisher Scientific) into a single cell suspension and plated at 15,000 cells/cm^2^ in E8 media with Y-27632 (10μM). Y-27632 was removed after 24 hours and cells cultured in E8 media until they reached approximately 70% confluency at which point the media was replaced with MelM media, this is considered day 0. On day 2 the media was replaced with PC/SMC1 media. On day 5 the media was replaced with PC/SMC2 media for 24 hours and then cells were passaged 1:3 on to fibronectin (FC01010MG, Fisher Scientific) coated tissue culture plates in SMC3 media. Plates were coated with fibronectin at 10ug/ml for 30 minutes at 37°C prior to the addition of cells. After 48 hours media was replaced with SMC4 media. After 48 hours the media was replaced with SMC5 and cells kept in culture for 7 days prior to freezing.

### Directed differentiation of hPSCs to brain microvascular endothelial cells

The differentiation of H1 hPSCs to brain microvascular endothelial cells was based on a previously published protocols^39, 69, 70^. Briefly, hPSCs were dissociated using Accutase (Thermo Fisher Scientific) into a single cell suspension and plated at 15,000 cells/cm^2^ in StemFlex media with Y-27632 (10μM). Y-27632 was removed after 24 hours and at 48 hours post-plating the media was replaced with Unconditioned Media (UM) (100ml Knock-out Serum Replacement (Thermo Fisher Scientific), 5ml non-essential ammino acids (Thermo Fisher Scientific), 2.5ml GlutaMax (Thermo Fisher Scientific), 5ml Pen/Strep (Thermo Fisher Scientific), 3.5μl β-mercapto-ethanol (Sigma), and 392.5ml DMEM/F12(1:1)), this was considered day zero. The media was replaced daily with fresh UM. On day six the media was changed to hESFM (Thermo Fisher Scientific) supplemented with 20 ng/mL bFGF (Peprotech), 10 µM retinoic acid (RA) (Sigma), and 1:50 B27 (Thermo Fisher Scientific). The media was not changed for 48 hours. On day eight, cells were washed once with DPBS and incubated with Accutase for 30 minutes at 37°C. Cells were collected via centrifugation and plated onto either standard tissue culture plates or transwell plates (Corning #3460). Both plates and transwells were coated with 400 µg/mL collagen IV (Sigma Aldrich) and 100 µg/mL fibronectin (Fisher Scientific) overnight with collagen and fibronectin and washed 1X with PBS prior to the addition of cells. For tissue culture plates, cells were plated at a density of 250,000 cells/cm^2^, whereas for transwell plates, cells were plated at a density of 1.1 x 10^6^ cells/cm^2^ on the transwell membrane. bFGF and RA were removed from the medium 24 hours after plating.

### Microvascular network formation

3D cell culture chips (idenTx, AIM Biotech) were utilized to generate *in vitro* microvascular networks (MVN)s. AIM chips contain three parallel channels: a central gel channel flanked by two media channels. Microposts separate fluidic channels and serve to confine the liquid gelling solution in the central channel by surface tension before polymerization. The gel channel is 1.3 mm wide and 0.25 mm tall, the gap between microposts is 0.1 mm, and the width of media channels is 0.5 mm. For vascular cell seeding, hPSC (iPS11)-derived ECs and PCs or SMCs were co-seeded into the chip as previously described^71^. Briefly, ECs and respective mural cells were concentrated in defined SFM media minus heparin containing thrombin (4 U/mL). Cell mixture solution was then further mixed with fibrinogen (3 mg/mL final concentration) at a 1:1 ratio and quickly pipetted into the chip through the gel inlet with a final concentration of 6 x10^6^/mL for ECs and 1.5 x10^6^/mL for mural cells. The device was placed upside down to polymerize in a humidified tip box and allowed to polymerize at 37 °C for 15 min in a 5% CO_2_ incubator, before endothelial serum free media was introduced to the media channels. After seeding, culture medium was added and changed daily in the device. After 7 days, MVNs are ready for further experiments.

### Cell characterization by bulk RNA sequencing and PCA

Primary HUVEC cells were obtained from VWR (Cat# 10171-906). Primary bronchial smooth muscle cells (BSMCs) were obtained from PromoCell (Cat# C12561). Primary pericytes were obtained from ScienCell (Cat#1200). RNA was extracted using the RNeasy Plus Mini kit (QIAGEN) following the manufacturer’s protocol. For bulk RNA sequencing of total RNA, libraries were prepared using the Swift RNA library prep kit (Swift Biosciences). RNA sequencing reads were mapped to the human genome (version GRCh38) using the STAR aligner (version 2.7.1a)^72^. The mapped reads were assigned to the human genes using featureCounts^73^ with the parameters ‘-p-s 2’, suitable for paired-end reversely-stranded libraries. Before performing PCA on the 5000 most variable genes, the normTransform function of DESeq2 (version 1.36.0)^74^ was used to normalize, center and log-transform (with a pseudocount of 1) the counts matrix. The ‘prcomp’ function in R (version 4.2.1) was used to carry out the PCA. PC2 and PC3 were used for computing Euclidean distances for the BSMC comparison, while PC1, PC2 and PC3 were used for the primary PC comparison. In the table of between-sample Euclidean distances (shown in Source Data file) each column is a comparison samples within two conditions. For computing pairwise Euclidean distances between pairs of conditions (Supplemental Fig. 1G-H) each group includes three replicates, so that there are 3 x 3 = 9 distance values corresponding to all possible pairs. For making a heatmap visualization of the marker gene expression, the normalized counts, *c_ij_*, for gene *i* in cell type *j* was standardized as *z_ij_* = (−µ*_i_*)/σ*_i_*.

### SARS-CoV-2 propagation, titration, and inactivation

SARS-CoV-2 (isolate USA_WA1/2020) was obtained from BEI Resources. BA.5.1 strain was obtained from a clinical isolate through the MassCPR Variant Program and verified by whole genome sequencing. The virus was propagated in Vero E6 cells (ATCC CRL-1586) cultured in Dulbecco’s modified Eagle’s medium (DMEM) supplemented with 2% fetal calf serum (FCS), penicillin (50 U/mL), and streptomycin (50 mg/mL). SARS-CoV-2 titer was determined in Vero E6 cells by plaque assay. work with SARS-CoV-2 was performed in the biosafety level 3 (BSL3) at the Ragon Institute (Cambridge, MA) following approved SOPs. To generate heat-inactivated SARS-CoV-2 a 1ml aliquot of SARS-CoV-2 was heated to 85°C for 15 minutes. For infection of SMCs, PCs, or ECs the cell culture media was removed and replaced with inoculating virus diluted in a minimum volume of culture media. The cells were placed at 37°C for 1 hour at which point the inoculum was removed and replaced with EC, PC, or SMC media or Infection media for conditioned media transfer experiments. For all conditioned media experiments control ECs were treated with infection media (unconditioned) for 48 hours.

### Bulk RNA-sequencing of infected and exposed vascular cells

ECs, SMCs or PCs were infected at an MOI of 1 or exposed to conditioned media. At the indicated time of collection RNA was extracted using the RNeasy Plus Mini kit (QIAGEN) following the manufacturer’s protocol. For bulk RNA sequencing of total RNA, libraries were prepared using the Swift RNA library prep kit (Swift Biosciences) or the SMARTerUltra-lowPOLYA-V4 (Takara Bio). Samples were sequenced on a NovaSeq 6000 sequencer.

### Analysis of gene expression

Paired-end reads (51x 51 bp) were mapped to a reference genome using the STAR aligner (v. 2.7.1a)^72^. The reference genome was a composite of the human genome (GRCh38) and that of the SARS-CoV-2 coronavirus (GenBank MN988713.1). Counts for human protein coding genes and lncRNAs (ENSEMBL release 93 annotations), together with the viral genes, were tabulated using featureCounts^73^. Differential expression of mRNAs was assessed for pairwise contrasts between conditions using estimated fold-changes and the Wald statistic in DESeq2 (v. 1.36.0)^74^. Per-gene dispersions were estimated using all conditions, as within-condition variance was observed to be relatively uniform across conditions in exploratory analyses. Unless otherwise noted, empirical Bayes shrinkage was applied to fold-change estimates^75^. Gene-set enrichment analysis was carried out between pairs of conditions using GSEA (v. 4.1.0)^76^ with counts that were normalized by DESeq2 and 1000 gene-set permutations were used to estimate p-values. Gene-sets that were tested for enrichment came from the Hallmark collection at the MSigDB^34^.

### Flow cytometry

HEK293T cells were obtained from ATCC and cultured in DMEM + 10% FBS and P/S, and split with 0.05% Trypsin for passaging. Human dermal fibroblasts (hDF) were obtained from Lonza and cultured in DMEM + 10% FBS and P/S, and split with 0.05% Trypsin for passaging. For live surface marker staining, cells were trypsinized with TrypLE Express for ∼5-7 or until cells detached. Cells were washed once with FACS staining buffer consisting of 2 mM EDTA and 0.5% BSA and centrifuged. Cells were then resuspended and incubated in FACS staining buffer and the respective fluorophore-conjugated antibodies for each cell type at the concentrations listed in Supplemental Table 1. For SMC staining that included intracellular SMA, cells were fixed with 1% PFA after the trypsinization and washing step listed above for 20 min at room temperature. The cells were then permeabilized in 90% methanol in PBS at −20C for at least 24 hr. After permeabilization, cells were washed in FACS buffer and resuspended in FACS buffer containing 0.1% Triton X-100 and the respective antibodies listed in Supplemental Table 1. All antibodies were done for 20-30 min at 4C and were followed by a wash with FACS buffer. Stained cells were analyzed on a BD Fortessa UV/Blue/Green/Red 4-laser flow cytometer. Cells were isolated as a population and then selected for singlets before setting gates to determine positive populations. Undifferentiated hPSCs served as biological controls for all stains.

### Immunofluorescence microscopy

Cells were fixed with 4% paraformaldehyde for 15 minutes at room temperature. Fixed cells were permeabilized for 15 minutes using 0.1% TX-100. Cells were then blocked with 3% BSA in PBS for 1 hour at room temperature. Cells were incubated with primary antibodies (see Supplemental Table 1) diluted in antibody buffer (0.01%TX-100 and 1%BSA in PBS) overnight at 4°C. Cells were washed with 0.05% Tween in PBS and incubated for 1 hour at room temperature with secondary antibodies (see Supplemental Table 1) diluted in antibody buffer. FITC-Phalloidin was added at 1:2000 and DAPI at 1:5000 with secondary antibodies.

### Imaging of reactive oxygen species production

Endothelial cells were exposed to SARS-CoV-2 for 48 hours. At 48 hours post-infection the media was removed and replaced with FluoroBrite media (A1896701, Thermo Fisher Scientific) supplemented with 1:50 B27 and CellRox Green Reagent (C10444, Thermo Fisher Scientific) at a final concentration of 5μM. Cells were incubated at 37°C for 30 minutes then washed 3 times with PBS and imaged immediately after washing. Cells were kept in FluoroBrite with B27 throughout imaging. The average intercellular florescence for five to seven cells for each condition was measured in each experiment. Three independent experiments were performed.

### SERPINE1 (PAI-1) and von Willebrand factor (vWF) release measurements

Endothelial cells (ECs) were exposed to media from SARS-CoV-2 infected or mock infected SMCs or PCs for 48 hours at which point the media was collected and analyzed for PAI-1 content using the Human Serpin E1/PAI-1 Quantikine ELISA Kit (DSE100, R&D Systems) according to the manufacturer’s protocol. The same media was analyzed for vWF content using the Human Von Willebrand Factor ELISA Kit (ab223864, Abcam) according to the manufacturer’s protocol. For analysis on the impact of SMC or PC treatment with IFNα or IFNγ cells were plated in a 24-well plate and treated with 100u/ml IFNα (ThermoFisher #111001) or 20ng/ml IFNγ(R&D 285-IF) in 300ul infection media for 24 hours at which point the cells were washed once with infection media and 300ul of fresh infection added and collected again 24 hours after SMC or PC conditioning. ECs were then treated with this conditioned media and the media was analyzed for vWF or SERPINE1 content as described above.

### Measurement of Tissue Factor Activity

At 48 hours post-infection SMCs were lysed and solubilized with 50 mM Tris-buffered saline (pH 8.0) containing 0.5% Triton X-100 or 0.1% Tween 20 at 4°C for 30 minutes. Samples were then centrifuged at 10,000xg for 10 minutes, and the supernatant was collected for analysis. Tissue factor (TF) activity was measured using a human tissue factor activity assay kit (ab108906, Abcam) according to the manufacturer’s protocol. Activity was normalized to total protein contents using the Pierce BCA Protein Assay Kit (FERA65453, Thermo Scientific).

### Measurement of transendothelial electrical resistance

BMECs were plated onto three Transwell filters (Corning #3460) using the procedure described in “Directed Differentiation of hPSCs to brain microvascular endothelial cells” ^69^. 48 hours after plating the media was removed from the apical side of the Transwell and replaced with SMC-conditioned media. At each time point, the TEER was measured in three separate wells. All data are represented as mean ± standard deviation for these collective measurements. Following the medium change on day 0 of subculture to remove bFGF and RA, no further medium changes were performed for the duration of the experiments.

### FITC-Dextran Permeability Assay

Endothelial cells were seeded in 12-well Transwell plates (Corning #3460) at 5e5 cells/Transwell. 72 hours after seeding ECs were either infected with SARS-CoV-2 (MOI 1) or treated with SMC conditioned media. 48 hours after treatment the media on the basolateral side was replaced with 1ml of fresh EC media. FITC-tagged dextran (10 μM) was suspended in 0.5 ml of EC medium and added to the apical side of the Transwell. To determine the level of transcytosis following 2 hours of incubation in a 37°C incubator on a rotating platform, we collected 150 μl from the 1ml of EC medium on the basolateral side of the Transwell. Fluorescence intensity in the collected media was measured using a SpectraMax iD3 plate reader set to 495-nm excitation and 519-nm emission settings.

### CellTiter Glo Viability Assay

Endothelial cells were plated in a 96 well plate at a density of 5e4 cells/well. 24 hours after plating the media was replaced with EC media containing 1uM or 5uM DMS or a matched volume of DMSO. 72 hours after treatment a CellTiter Glo 2.0 (Promega # G9241) assay was performed according the manufacturer instructions.

## Data availability

The RNA sequencing data generated in this study has been deposited in the GEO database under accession code GSE240766. All additional data generated in this study are provided in the Supplementary Information/Source Data file.

## Acknowledgments

We acknowledge funding from the Wellcome-LEAP HOPE initiative, NIH (T32 EB016652), the Wyss Institute, and the Whitehead Institute. We thank the Genome Technology Core and Keck Imaging Facility at Whitehead Institute for Biomedical Research. This work was supported by NIH grant U19AI131135, NIBIB grant T32 EB016652, and by a generous gift from Jim Stone. The following reagent was deposited by the Centers for Disease Control and Prevention and obtained through BEI Resources, NIAID, NIH: SARS-Related Coronavirus 2, Isolate USA-WA1/2020, NR-52281. This work was performed in part in the Ragon Institute BSL3 core, which is supported by the NIH-funded Harvard University Center for AIDS Research (P30 AI060354) and the Massachusetts Consortium on Pathogen Readiness (MassCPR).

## Author Contributions

A.R., A.K., R.J., and D.M designed the project. T.W. and X.G. performed the bulk RNA sequencing data analysis. T.L. assisted in stem cell maintenance and differentiations. R.K. and Z.W. generated and analyzed microfluidic cultures of hPSC-derived vascular cells. M.F. assisted with real-time PCR data collection. L.G. facilitated work in the Ragon BSL3 facility. A.R. and A.K. analyzed all data except for RNA sequencing data and prepared the manuscript. All authors discussed the results and commented on the manuscript.

## References

1. Arora, P., Jafferany, M., Lotti, T., Sadoughifar, R. & Goldust, M. Learning from history: Coronavirus outbreaks in the past. Dermatol Ther 33, e13343 (2020).

2. WHO, Vol. 2021 (

3. Acharya, Y., Alameer, A., Calpin, G., Alkhattab, M. & Sultan, S. A comprehensive review of vascular complications in COVID-19. J Thromb Thrombolysis 53, 586–593 (2022).

4. Merschel, M. in American Heart Association News Stories, Edn. February 24, 2023 (2023).

5. Xie, Y., Xu, E., Bowe, B. & Al-Aly, Z. Long-term cardiovascular outcomes of COVID-19. Nat Med 28, 583–590 (2022).

6. Spyropoulos, A.C., et al. Occurrence of Thromboembolic Events and Mortality Among Hospitalized Coronavirus 2019 Patients: Large Observational Cohort Study of Electronic Health Records. TH Open 6, e408–e420 (2022).

7. Belani, P. et al. COVID-19 Is an Independent Risk Factor for Acute Ischemic Stroke. AJNR Am J Neuroradiol 41, 1361–1364 (2020).

8. Ackermann, M. et al. Pulmonary Vascular Endothelialitis, Thrombosis, and Angiogenesis in Covid-19. N Engl J Med 383, 120–128 (2020).

9. Schimmel, L. et al. Endothelial cells are not productively infected by SARS-CoV-2. Clin Transl Immunology 10, e1350 (2021).

10. Urata, R. et al. Senescent endothelial cells are predisposed to SARS-CoV-2 infection and subsequent endothelial dysfunction. Sci Rep 12, 11855 (2022).

11. McQuaid, C. & Montagne, A. SARS-CoV-2 and vascular dysfunction: a growing role for pericytes. Cardiovasc Res (2022).

12. Stein, S.R. et al. SARS-CoV-2 infection and persistence in the human body and brain at autopsy. Nature 612, 758–763 (2022).

13. Ryu, G. & Shin, H.W. SARS-CoV-2 Infection of Airway Epithelial Cells. Immune Netw 21, e3 (2021).

14. Varga, Z. et al. Endothelial cell infection and endotheliitis in COVID-19. Lancet 395, 1417–1418 (2020).

15. Merad, M. & Martin, J.C. Pathological inflammation in patients with COVID-19: a key role for monocytes and macrophages. Nat Rev Immunol 20, 355–362 (2020).

16. Blanco-Melo, D. et al. Imbalanced Host Response to SARS-CoV-2 Drives Development of COVID-19. Cell 181, 1036–1045 e1039 (2020).

17. Lilly, B. We have contact: endothelial cell-smooth muscle cell interactions. Physiology (Bethesda) 29, 234–241 (2014).

18. Mendez-Barbero, N., Gutierrez-Munoz, C. & Blanco-Colio, L.M. Cellular Crosstalk between Endothelial and Smooth Muscle Cells in Vascular Wall Remodeling. Int J Mol Sci 22 (2021).

19. Patsch, C. et al. Generation of vascular endothelial and smooth muscle cells from human pluripotent stem cells. Nat Cell Biol 17, 994–1003 (2015).

20. Kumar, A. et al. Specification and Diversification of Pericytes and Smooth Muscle Cells from Mesenchymoangioblasts. Cell Rep 19, 1902–1916 (2017).

21. Koyano-Nakagawa, N. & Garry, D.J. Etv2 as an essential regulator of mesodermal lineage development. Cardiovasc Res 113, 1294–1306 (2017).

22. Cao, G. et al. How vascular smooth muscle cell phenotype switching contributes to vascular disease. Cell Commun Signal 20, 180 (2022).

23. Wanjare, M., Kuo, F. & Gerecht, S. Derivation and maturation of synthetic and contractile vascular smooth muscle cells from human pluripotent stem cells. Cardiovasc Res 97, 321–330 (2013).

24. Sola, I., Almazan, F., Zuniga, S. & Enjuanes, L. Continuous and Discontinuous RNA Synthesis in Coronaviruses. Annu Rev Virol 2, 265–288 (2015).

25. Hoffmann, M. et al. SARS-CoV-2 Cell Entry Depends on ACE2 and TMPRSS2 and Is Blocked by a Clinically Proven Protease Inhibitor. Cell 181, 271–280 e278 (2020).

26. Cantuti-Castelvetri, L. et al. Neuropilin-1 facilitates SARS-CoV-2 cell entry and infectivity. Science 370, 856–860 (2020).

27. Fantin, A. et al. NRP1 acts cell autonomously in endothelium to promote tip cell function during sprouting angiogenesis. Blood 121, 2352–2362 (2013).

28. Lei, Y. et al. SARS-CoV-2 Spike Protein Impairs Endothelial Function via Downregulation of ACE 2. Circ Res 128, 1323–1326 (2021).

29. Wolter, N. et al. Early assessment of the clinical severity of the SARS-CoV-2 omicron variant in South Africa: a data linkage study. Lancet 399, 437–446 (2022).

30. Loveday, E.K. et al. Effect of Inactivation Methods on SARS-CoV-2 Virion Protein and Structure. Viruses 13 (2021).

31. Boueiz, A. & Hassoun, P.M. Regulation of endothelial barrier function by reactive oxygen and nitrogen species. Microvasc Res 77, 26–34 (2009).

32. Ma, J., Sanchez-Duffhues, G., Goumans, M.J. & Ten Dijke, P. TGF-beta-Induced Endothelial to Mesenchymal Transition in Disease and Tissue Engineering. Front Cell Dev Biol 8, 260 (2020).

33. Cooley, B.C. et al. TGF-beta signaling mediates endothelial-to-mesenchymal transition (EndMT) during vein graft remodeling. Sci Transl Med 6, 227ra234 (2014).

34. Liberzon, A. et al. The Molecular Signatures Database (MSigDB) hallmark gene set collection. Cell Syst 1, 417–425 (2015).

35. Lopez-Castaneda, S. et al. Inflammatory and Prothrombotic Biomarkers Associated With the Severity of COVID-19 Infection. Clin Appl Thromb Hemost 27, 1076029621999099 (2021).

36. Choudhary, S., Sharma, K. & Singh, P.K. Von Willebrand factor: A key glycoprotein involved in thrombo-inflammatory complications of COVID-19. Chem Biol Interact 348, 109657 (2021).

37. Helms, J. et al. High risk of thrombosis in patients with severe SARS-CoV-2 infection: a multicenter prospective cohort study. Intensive Care Med 46, 1089–1098 (2020).

38. Greene, C. et al. Author Correction: Blood-brain barrier disruption and sustained systemic inflammation in individuals with long COVID-associated cognitive impairment. Nat Neurosci 27, 1019 (2024).

39. Stebbins, M.J. et al. Differentiation and characterization of human pluripotent stem cell-derived brain microvascular endothelial cells. Methods 101, 93–102 (2016).

40. Helms, H.C. et al. In vitro models of the blood-brain barrier: An overview of commonly used brain endothelial cell culture models and guidelines for their use. J Cereb Blood Flow Metab 36, 862–890 (2016).

41. Yang, R.C. et al. SARS-CoV-2 productively infects human brain microvascular endothelial cells. J Neuroinflammation 19, 149 (2022).

42. Srinivasan, B. et al. TEER measurement techniques for in vitro barrier model systems. J Lab Autom 20, 107–126 (2015).

43. Wang, L. et al. Vascular smooth muscle-derived tissue factor is critical for arterial thrombosis after ferric chloride-induced injury. Blood 113, 705–713 (2009).

44. Nguyen, D., Jeon, H.M. & Lee, J. Tissue factor links inflammation, thrombosis, and senescence in COVID-19. Sci Rep 12, 19842 (2022).

45. Subrahmanian, S. et al. Tissue factor upregulation is associated with SARS-CoV-2 in the lungs of COVID-19 patients. J Thromb Haemost 19, 2268–2274 (2021).

46. Spiegel, S. & Milstien, S. The outs and the ins of sphingosine-1-phosphate in immunity. Nat Rev Immunol 11, 403–415 (2011).

47. Yanagida, K. & Hla, T. Vascular and Immunobiology of the Circulatory Sphingosine 1-Phosphate Gradient. Annu Rev Physiol 79, 67–91 (2017).

48. Schuchardt, M., Tolle, M., Prufer, J. & van der Giet, M. Pharmacological relevance and potential of sphingosine 1-phosphate in the vascular system. Br J Pharmacol 163, 1140–1162 (2011).

49. McGowan, E.M., Haddadi, N., Nassif, N.T. & Lin, Y. Targeting the SphK-S1P-SIPR Pathway as a Potential Therapeutic Approach for COVID-19. Int J Mol Sci 21 (2020).

50. Vanlandewijck, M. et al. A molecular atlas of cell types and zonation in the brain vasculature. Nature 554, 475–480 (2018).

51. Armulik, A., Genove, G. & Betsholtz, C. Pericytes: developmental, physiological, and pathological perspectives, problems, and promises. Dev Cell 21, 193–215 (2011).

52. von Tell, D., Armulik, A. & Betsholtz, C. Pericytes and vascular stability. Exp Cell Res 312, 623–629 (2006).

53. Niu, X. et al. EDIL3 influenced the alphavbeta3-FAK/MEK/ERK axis of endothelial cells in psoriasis. J Cell Mol Med 26, 5202–5212 (2022).

54. Gajate-Arenas, M. et al. The Immune Response of OAS1, IRF9, and IFI6 Genes in the Pathogenesis of COVID-19. Int J Mol Sci 25 (2024).

55. Bizzotto, J. et al. SARS-CoV-2 Infection Boosts MX1 Antiviral Effector in COVID-19 Patients. iScience 23, 101585 (2020).

56. Nain, Z. et al. Transcriptomic studies revealed pathophysiological impact of COVID-19 to predominant health conditions. Brief Bioinform 22 (2021).

57. Parker, N. & Porter, A.C. Identification of a novel gene family that includes the interferon-inducible human genes 6-16 and ISG12. BMC Genomics 5, 8 (2004).

58. Mackelprang, R.D. et al. Upregulation of IFN-stimulated genes persists beyond the transitory broad immunologic changes of acute HIV-1 infection. iScience 26, 106454 (2023).

59. Fusco, D.N. et al. HELZ2 Is an IFN Effector Mediating Suppression of Dengue Virus. Front Microbiol 8, 240 (2017).

60. Bocci, M. et al. Infection of Brain Pericytes Underlying Neuropathology of COVID-19 Patients. Int J Mol Sci 22 (2021).

61. Khan, A.O. et al. Preferential uptake of SARS-CoV-2 by pericytes potentiates vascular damage and permeability in an organoid model of the microvasculature. Cardiovasc Res 118, 3085–3096 (2022).

62. Xu, S.W., Ilyas, I. & Weng, J.P. Endothelial dysfunction in COVID-19: an overview of evidence, biomarkers, mechanisms and potential therapies. Acta Pharmacol Sin 44, 695–709 (2023).

63. Rauti, R. et al. Effect of SARS-CoV-2 proteins on vascular permeability. Elife 10 (2021).

64. Li, Y. et al. SARS-CoV-2 induces double-stranded RNA-mediated innate immune responses in respiratory epithelial-derived cells and cardiomyocytes. Proc Natl Acad Sci U S A 118 (2021).

65. Girgis, S. et al. Evolution of naturally arising SARS-CoV-2 defective interfering particles. Commun Biol 5, 1140 (2022).

66. Al-Samkari, H. et al. COVID-19 and coagulation: bleeding and thrombotic manifestations of SARS-CoV-2 infection. Blood 136, 489–500 (2020).

67. Xu, S.W., Ilyas, I. & Weng, J.P. Endothelial dysfunction in COVID-19: an overview of evidence, biomarkers, mechanisms and potential therapies. Acta Pharmacol Sin, 1–15 (2022).

68. Jiang, Z. et al. NOX2 and NOX5 are increased in cardiac microvascular endothelium of deceased COVID-19 patients. Int J Cardiol 370, 454–462 (2023).

69. Qian, T. et al. Directed differentiation of human pluripotent stem cells to blood-brain barrier endothelial cells. Sci Adv 3, e1701679 (2017).

70. Neal, E.H. et al. A Simplified, Fully Defined Differentiation Scheme for Producing Blood-Brain Barrier Endothelial Cells from Human iPSCs. Stem Cell Reports 12, 1380–1388 (2019).

71. Wan, Z. et al. A robust vasculogenic microfluidic model using human immortalized endothelial cells and Thy1 positive fibroblasts. Biomaterials 276, 121032 (2021).

72. Dobin, A. et al. STAR: ultrafast universal RNA-seq aligner. Bioinformatics 29, 15–21 (2013).

73. Liao, Y., Smyth, G.K. & Shi, W. featureCounts: an efficient general purpose program for assigning sequence reads to genomic features. Bioinformatics 30, 923–930 (2014).

74. Love, M.I., Huber, W. & Anders, S. Moderated estimation of fold change and dispersion for RNA-seq data with DESeq2. Genome Biol 15, 550 (2014).

75. Stephens, M. False discovery rates: a new deal. Biostatistics 18, 275–294 (2017).

76. Subramanian, A. et al. Gene set enrichment analysis: a knowledge-based approach for interpreting genome-wide expression profiles. Proc Natl Acad Sci U S A 102, 15545–15550 (2005).

77. Finkel, Y. et al. The coding capacity of SARS-CoV-2. Nature 589, 125–130 (2021).

