## Supplemental Figures and Tables for "SARS-CoV-2 infection of human pluripotent stem cell-derived vascular cells reveals smooth muscle cells as key mediators of vascular pathology during infection"

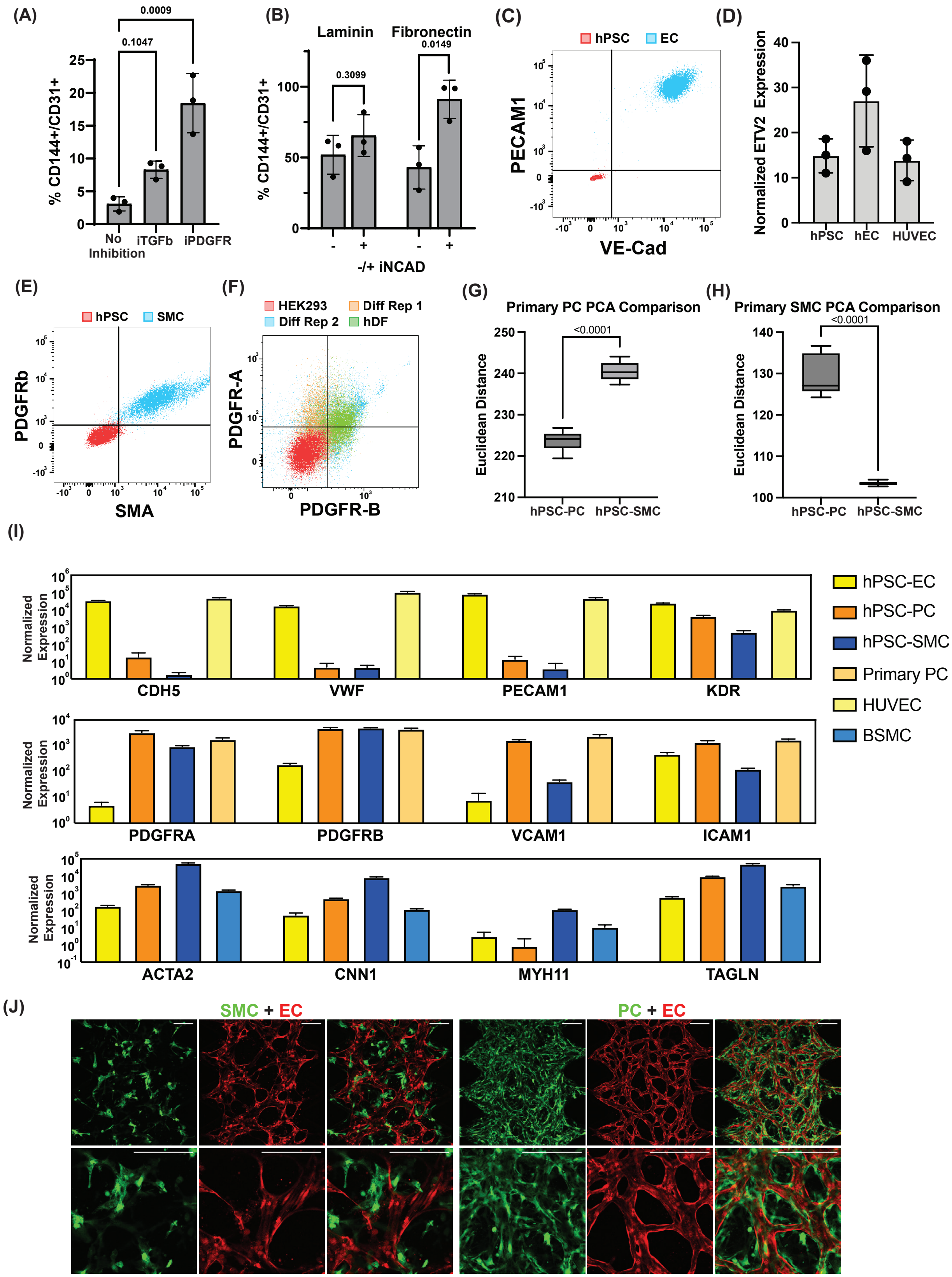
(A)
0.0009
0.1047
% CD144+/CD31+
No Inhibition
iTGFb
iPDGFR
(B)
Laminin
Fibronectin
0.3099
0.0149
% CD144+/CD31+
-/+ iNCAD
(C)
hPSC
EC
PECAM1
VE-Cad
(D)
Normalized ETV2 Expression
hPSC
hEC
HUVEC
(E)
hPSC
SMC
PDGFRb
SMA
(F)
HEK293
Diff Rep 1
Diff Rep 2
hDF
PDGFR-A
PDGFR-B
(G)
Primary PC PCA Comparison
<0.0001
Euclidean Distance
hPSC-PC
hPSC-SMC
(H)
Primary SMC PCA Comparison
<0.0001
Euclidean Distance
hPSC-PC
hPSC-SMC
(I)
Normalized Expression
CDH5
VWF
PECAM1
KDR
PDGFRA
PDGFRB
VCAM1
ICAM1
ACTA2
CNN1
MYH11
TAGLN
hPSC-EC
hPSC-PC
hPSC-SMC
Primary PC
HUVEC
BSMC
(J)
SMC + EC
PC + EC

**Supplemental Fig. 1: Characterization of hPSC-derived endothelial cells, pericytes, and smooth muscle cells.** (A) Quantification of EC derivation efficiency by flow cytometry of double-positive PECAM1 and VE-Cad on two replicate differentiations with the dependence of either TGFβ or PDGFR inhibition. Bar graph shows mean value and error bar shows +/- SD. Conditions were compared using a one-way ANOVA with Dunnett's multiple comparisons test, with a single pooled variance (B) Quantification of EC derivation efficiency by flow cytometry of double-positive PECAM1 and VE-Cad with the dependence of passaging on fibronectin or laminin and with or without N-cadherin inhibition (+/- iNCAD). Bar graph shows mean value and error bar shows +/- SD. Experimental conditions were compared using multiple unpaired t tests with the two-stage linear step-up procedure of Benjamini, Krieger, and Yekutieli (BKY). (C) Representative flow cytometry dot plot of double-positive PECAM1 and VE-Cad expression after purification, expansion, cryopreservation, and thawing of hPSC-derived ECs. (D) Normalized ETV2 RNA expression in hSPC-derived ECs, hPSCs, and primary HUVECs (E) Representative flow cytometry dot plot of double-positive PDGFRb and SMA expression after purification, expansion, cryopreservation, and thawing of hPSC-derived SMCs. (F) Representative flow cytometry dot plot of double-positive PDGFR-B and PDGFR-A expression after purification, expansion, cryopreservation, and thawing of hPSC-derived PCs. (G)Euclidean distance between Primary PCs and hPSC-derived PCs or hPSC-derived SMCs. Center line represents the median value of 240.3 for hPSC-SMC comparison and 224.1 for hPSC-PC comparison. (H) Euclidean distance between Primary BSMCs and hPSC-derived PCs or hPSC-derived SMCs. Center line represents the median value of 103.4 for hPSC-SMC comparison and 127.1 for hPSC-PC comparison (I) Normalized ENA expression of EC markers genes (top), PC marker genes (middle), and SMC marker genes (bottom). Values are average expression levels for three separate differentiations of all hPSC-derived cells. For primary cells (HUVEC, BSMC, and Primary PC) values are average expression from RNA isolated from three separate cultures. (J) Images of mSCarlett-tagged hPSC derived ECs and GFP-tagged hPSC-derived SMCs or PC seeded in microfluidic chambers. The experiment was performed twice with similar results. Representative images from a single experiment are shown.

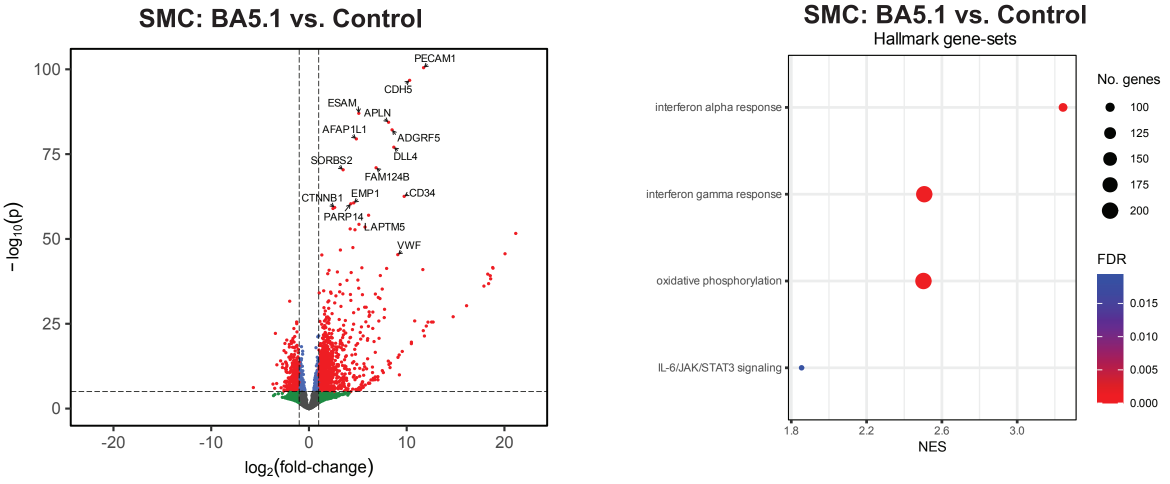


**Supplemental Fig. 2: Exposure of SMCs to the Omicron (BA5.1) variant activates inflammatory signaling.** hPSC-derived SMCs were infected with the Omicron variant (BA5.1) at an MOI of 1. Bulk RNA sequencing was performed on RNA isolated from hPSC-derived SMCs 48 hours after virus exposure. Volcano plots showing differential gene expression compared to control uninfected SMCs. Gene set enrichment analysis (GSEA)[76] was performed on differentially expressed genes to analyze the transcriptional response to infection. Dot plots show gene-sets from the Hallmark collection[34] of the MSigDB that were enriched (FDR < 0.05).

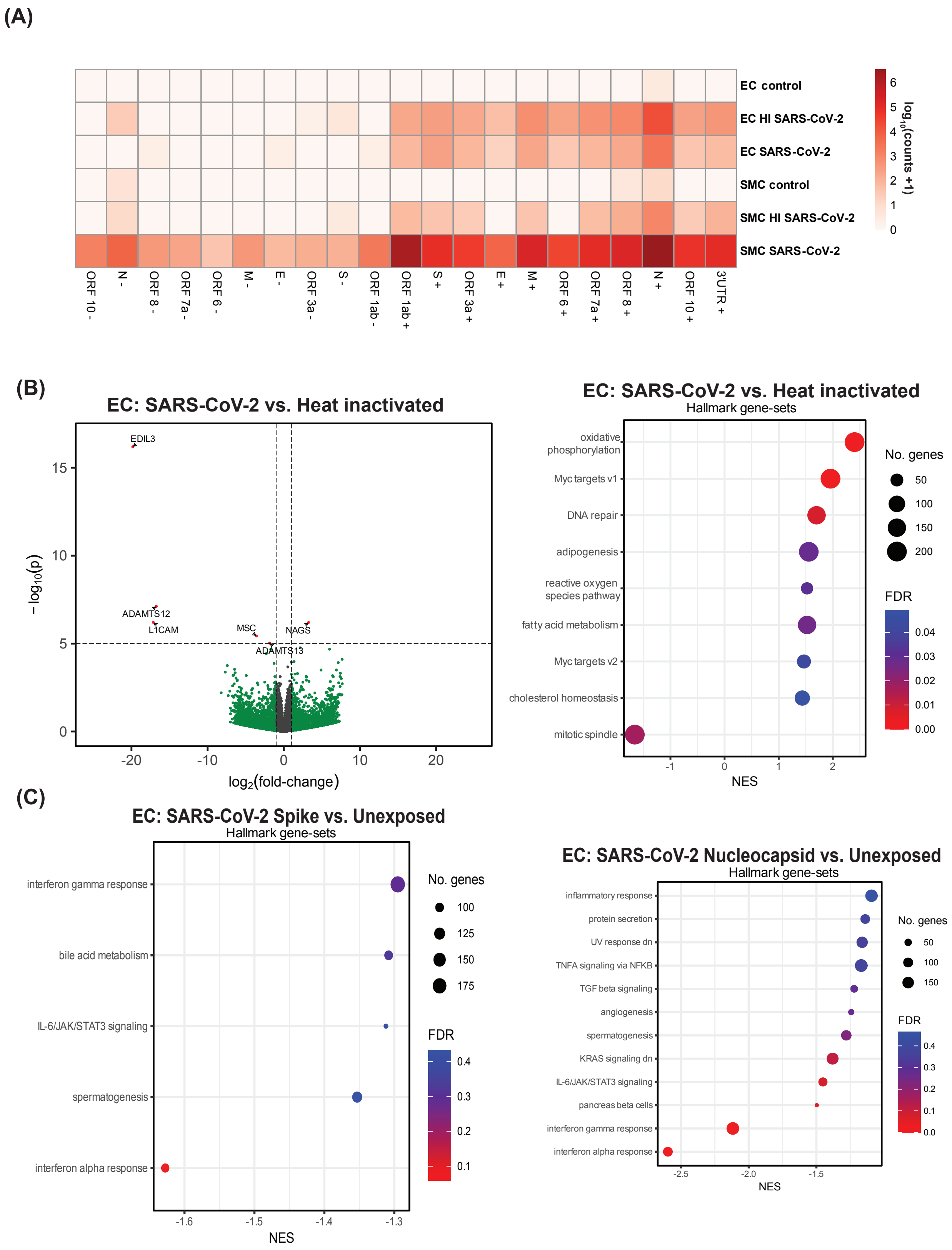
(A)
EC control
EC HI SARS-CoV-2
EC SARS-CoV-2
SMC control
SMC HI SARS-CoV-2
SMC SARS-CoV-2
ORF 10 -
N -
ORF 8 -
ORF 7a -
ORF 6 -
M -
E -
ORF 3a -
S -
ORF 1ab -
ORF 1ab +
S +
ORF 3a +
E +
M +
ORF 6 +
ORF 7a +
ORF 8 +
N +
ORF 10 +
3UTR +
$\log_{10}$(counts +1)
0
1
2
3
4
5
6
(B)
EC: SARS-CoV-2 vs. Heat inactivated
EDIL3
ADAMTS12
L1CAM
MSC
NAGS
ADAMTS13
$-\log_{10}(p)$
$\log_2$(fold-change)
EC: SARS-CoV-2 vs. Heat inactivated
Hallmark gene-sets
oxidative phosphorylation
Myc targets v1
DNA repair
adipogenesis
reactive oxygen species pathway
fatty acid metabolism
Myc targets v2
cholesterol homeostasis
mitotic spindle
NES
No. genes
50
100
150
200
FDR
0.04
0.03
0.02
0.01
0.00
(C)
EC: SARS-CoV-2 Spike vs. Unexposed
Hallmark gene-sets
interferon gamma response
bile acid metabolism
IL-6/JAK/STAT3 signaling
spermatogenesis
interferon alpha response
NES
No. genes
100
125
150
175
FDR
0.4
0.3
0.2
0.1
EC: SARS-CoV-2 Nucleocapsid vs. Unexposed
Hallmark gene-sets
inflammatory response
protein secretion
UV response dn
TNFA signaling via NFKB
TGF beta signaling
angiogenesis
spermatogenesis
KRAS signaling dn
IL-6/JAK/STAT3 signaling
pancreas beta cells
interferon gamma response
interferon alpha response
NES
No. genes
50
100
150
FDR
0.4
0.3
0.2
0.1
0.0

**Supplemental Fig. 3: EC exposure to SARS-CoV-2 results in induction of metabolic and reactive oxygen species pathways.** (A) Heatmap representation of mRNA levels for the ORFs of SARS-CoV-2[77] in each sample. The counts are visualized as $\log_{10}$(counts + 1) where the counts are DESeq2 normalized counts. Values shown are the averaged value of two independent experiments. (+) indicates positive sense mRNA, (-) indicates negative sense mRNA. (B) hPSC derived ECs were exposed to live SARS-CoV-2 (MOI=1) or heat-inactivated SARS-CoV-2. Bulk RNA sequencing was performed on RNA isolated at 48 hours post exposure. Volcano plots showing differential gene expression ECs exposed to live SARS-CoV-2 vs heat-inactivated SARS-CoV-2. Dot plots show gene-sets from the Hallmark collection[34] of the MSigDB that were enriched (FDR < 0.05) using gene-set enrichment analysis (GSEA)[76]. A full list of differentially expressed genes can be found in the Source Data file. (C) Bulk RNA sequencing was performed on RNA isolated from hPSC-derived ECs 48 hours after exposing cells to purified SARS-CoV-2 spike or nucleocapsid proteins. Gene set enrichment analysis was performed on sequenced samples. Dot plots show gene-sets from the Hallmark collection[34].

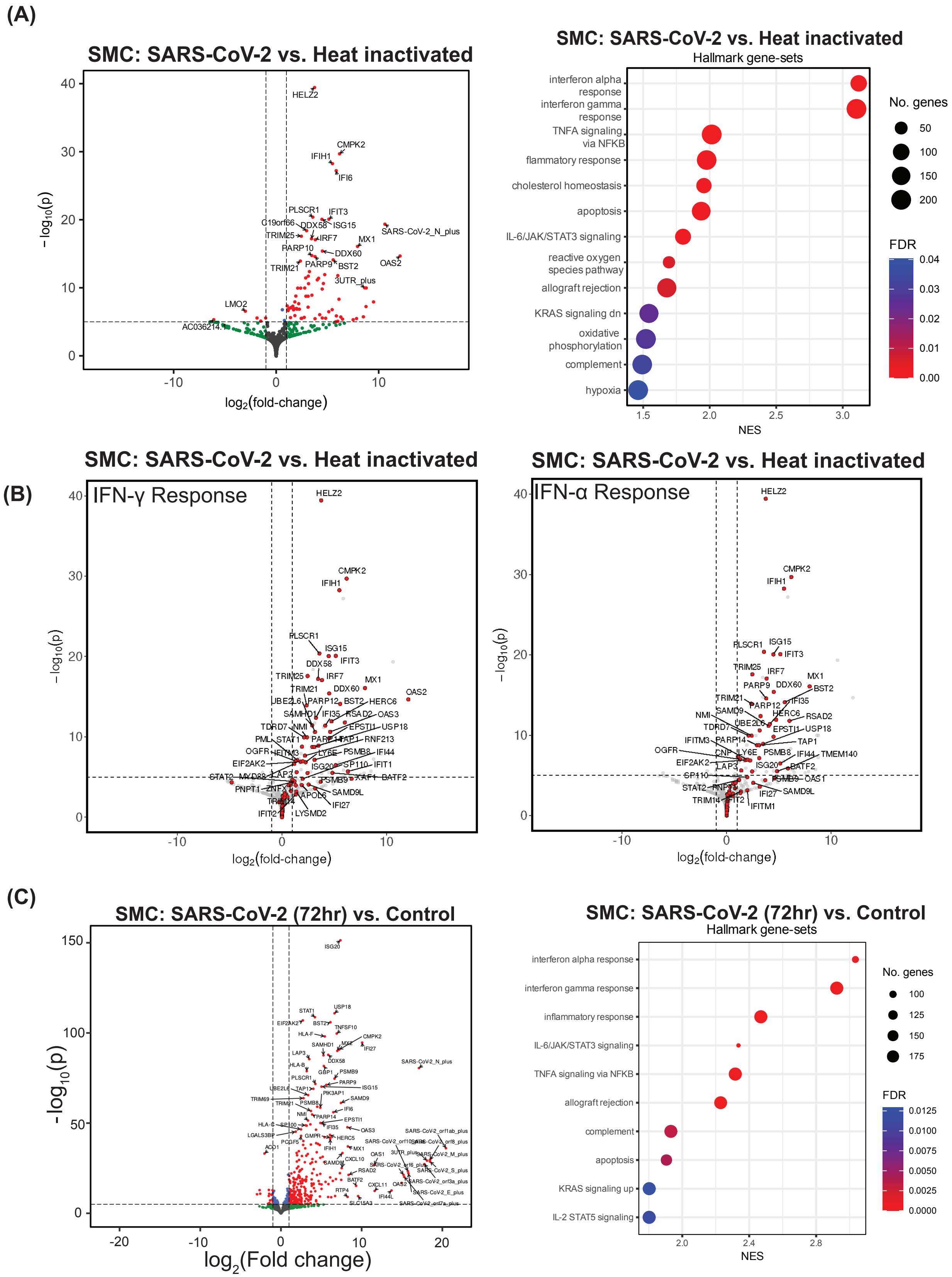
(A)
SMC: SARS-CoV-2 vs. Heat inactivated
-log10(p)
log2(fold-change)
HELZ2
CMPK2
IFIH1
IFI6
PLSCR1
IFIT3
C19orf66
DDX58
ISG15
TRIM25
IRF7
SARS-CoV-2_N_plus
MX1
PARP10
DDX60
PARP9
BST2
OAS2
TRIM21
3UTR_plus
LMO2
AC036214.1
SMC: SARS-CoV-2 vs. Heat inactivated
Hallmark gene-sets
interferon alpha response
interferon gamma response
TNFA signaling via NFKB
flammatory response
cholesterol homeostasis
apoptosis
IL-6/JAK/STAT3 signaling
reactive oxygen species pathway
allograft rejection
KRAS signaling dn
oxidative phosphorylation
complement
hypoxia
NES
No. genes
FDR
(B)
SMC: SARS-CoV-2 vs. Heat inactivated
IFN-γ Response
-log10(p)
log2(fold-change)
SMC: SARS-CoV-2 vs. Heat inactivated
IFN-α Response
-log10(p)
log2(fold-change)
(C)
SMC: SARS-CoV-2 (72hr) vs. Control
-log10(p)
log2(Fold change)
ISG20
SMC: SARS-CoV-2 (72hr) vs. Control
Hallmark gene-sets
interferon alpha response
interferon gamma response
inflammatory response
IL-6/JAK/STAT3 signaling
TNFA signaling via NFKB
allograft rejection
complement
apoptosis
KRAS signaling up
IL-2 STAT5 signaling
NES
No. genes
FDR

**Supplemental Fig. 4: Productive SARS-CoV-2 infection results in amplification of inflammatory signaling in hPSC-derived SMCs.** The transcriptional responses of hPSC-derived SMCs following addition of live SARS-CoV-2 (MOI=1) or an equal volume of heat-inactivated SARS-CoV-2 were examined by bulk-RNA sequencing 48 hours after virus exposure. A full list of differentially expressed genes can be found in the Source Data file. **(A)** Volcano plots showing differential gene expression in SMCs exposed to live SARS-CoV-2 or heat-inactivated SARS-CoV-2. Dot plots show gene-sets from the Hallmark collection[34] of the MSigDB that were enriched (FDR < 0.05) using gene-set enrichment analysis (GSEA)[76] **(B)** The volcano plots show genes that are differentially expressed in SMCs exposed to live SARS-CoV-2 compared to SMCs exposed to heat-inactivated SARS-CoV-2 and highlight the "IFN-γ response" (left) and "IFN-α response" (right) gene-sets from the Hallmark collection[34] of the MSigDB. Genes that are highlighted in red belong to the respective interferon response gene-set, with GSEA "leading edge" genes labeled by name. **(C)** Bulk RNA sequencing was performed on RNA isolated from hPSC-derived SMCs 72 hours after exposure to live SARS-CoV-2 (MOI=1). Volcano plots showing differential gene expression compared to control uninfected SMCs. Gene set enrichment analysis (GSEA)[76] was performed on differentially expressed genes to analyze the transcriptional response to infection. Dot plots show gene-sets from the Hallmark collection[34] of the MSigDB that were enriched (FDR < 0.05).

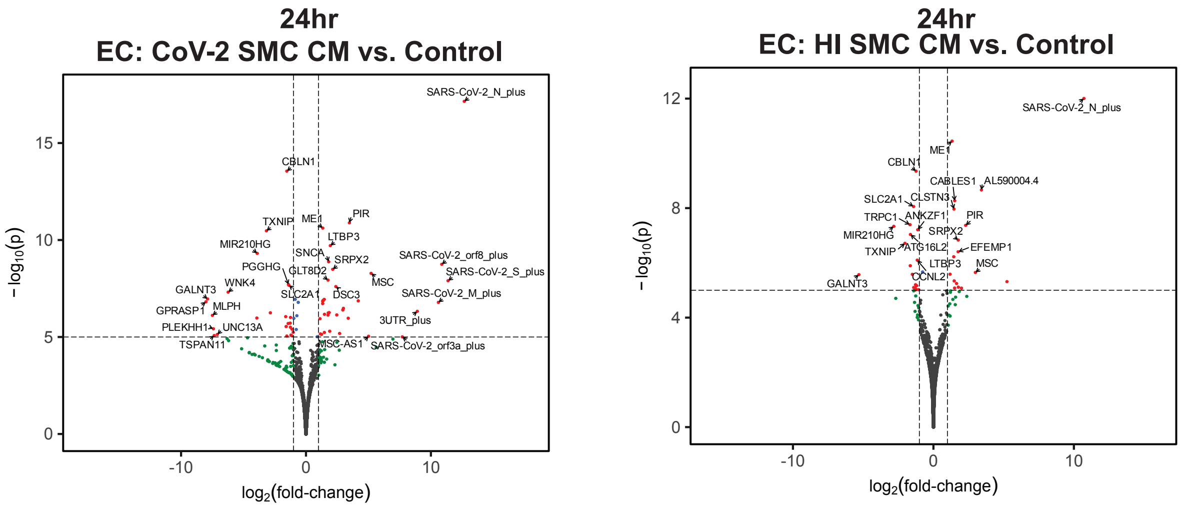


**Supplemental Fig. 5: Early response of ECs to factors secreted from SARS-CoV-2 exposed SMCs:** Bulk RNA sequencing was performed on RNA isolated from ECs 24 hours after exposure to SMC-conditioned media (Figure 4A). The volcano plots show genes that are differentially expressed in ECs exposed to media from SARS-CoV-2 infected SMCs (*CoV-2 SMC CM*) (left) compared to control ECs (*Control)* and ECs exposed to media from SMCs exposed to heat-inactivated SARS-CoV-2 *(HI SMC CM)* compared to control ECs *(Control) (right)*. A full list of differentially expressed genes can be found in the Source Data file.

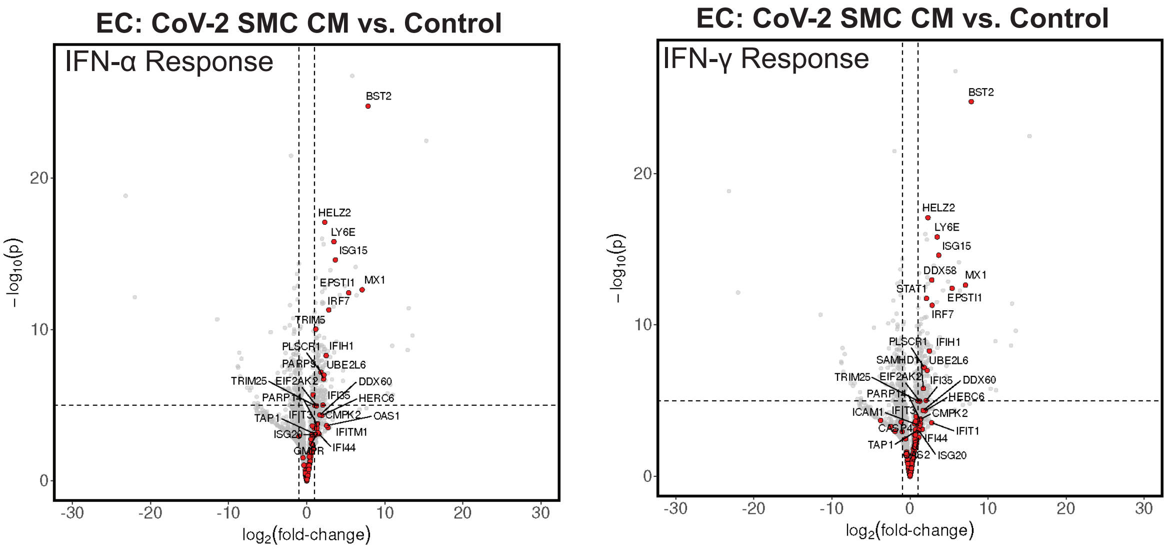


**Supplemental Fig. 6: SARS-CoV-2 infected SMCs release factors that promote inflammatory signaling in ECs.** Bulk RNA sequencing was performed on RNA isolated from ECs 48 hours after exposure to SMC-conditioned media **(**Figure 4A) as well control ECs. Genes from the “IFN-αresponse” (left) and “IFN-γ response” (right) gene-sets of the MSigDB’s Hallmark collection[34] are highlighted in red, with GSEA “leading edge” genes indicated by name.

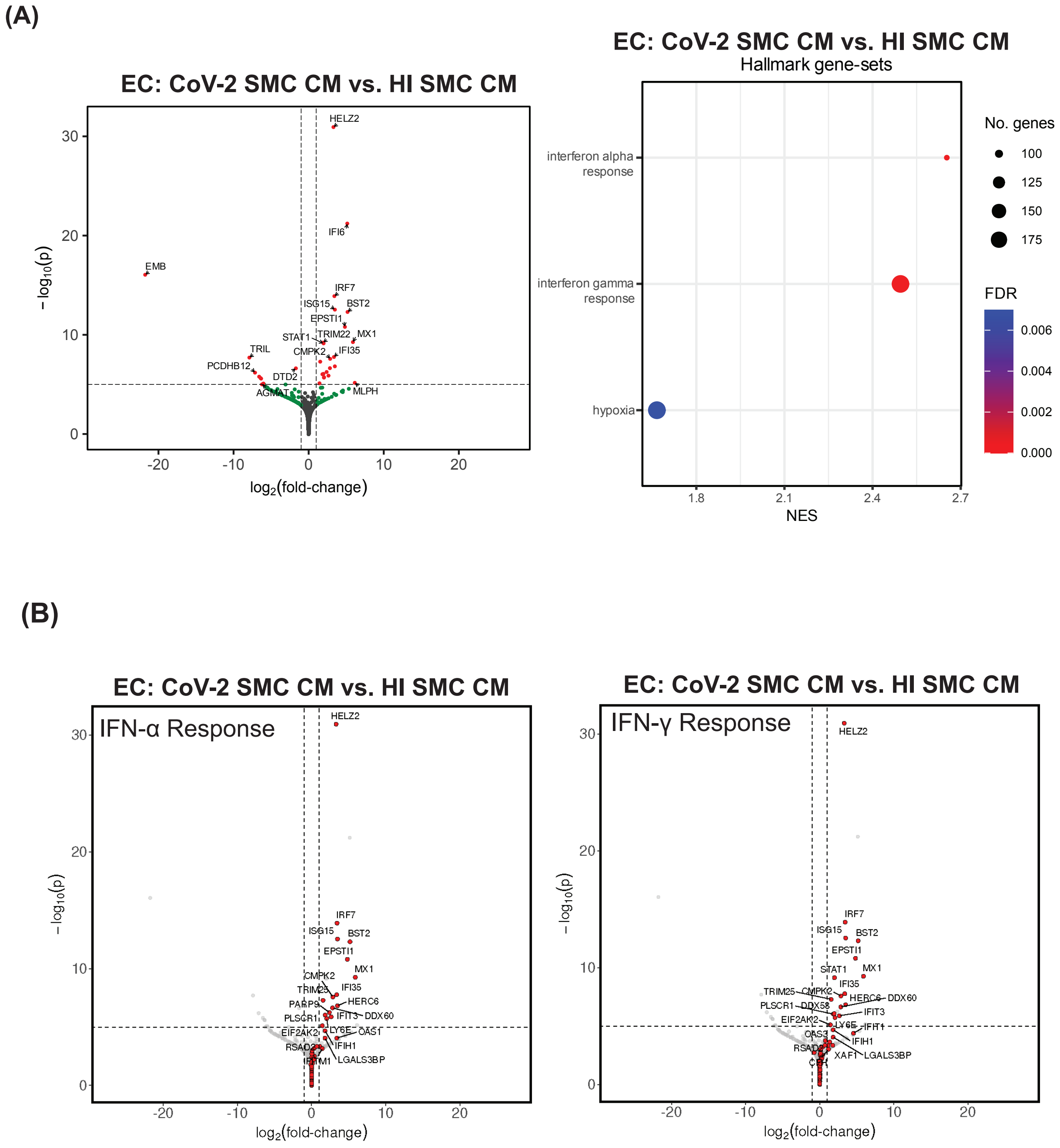


**Supplemental Fig 7: SARS-CoV-2 infection of SMCs amplifies the release of factors the promote inflammatory signaling in ECs.** Bulk RNA sequencing was performed on RNA isolated from ECs 48 hours after exposure to media from SMCs infected with SARS-CoV-2 or exposed to media from SMCs treated with heat-inactivated SARS-CoV-2 **(A)** Volcano plots showing differential gene expression. Dot plots show gene-sets from the Hallmark collection[34] of the MSigDB that were enriched (FDR < 0.05) using gene-set enrichment analysis (GSEA) **(B)** The volcano plots highlight the “IFN-α response” (left) and “IFN-γ response” (right) gene-sets from the Hallmark collection[34] of the MSigDB. Genes that are highlighted in red belong to the respective interferon response gene-set, with GSEA “leading edge” genes labeled by name.

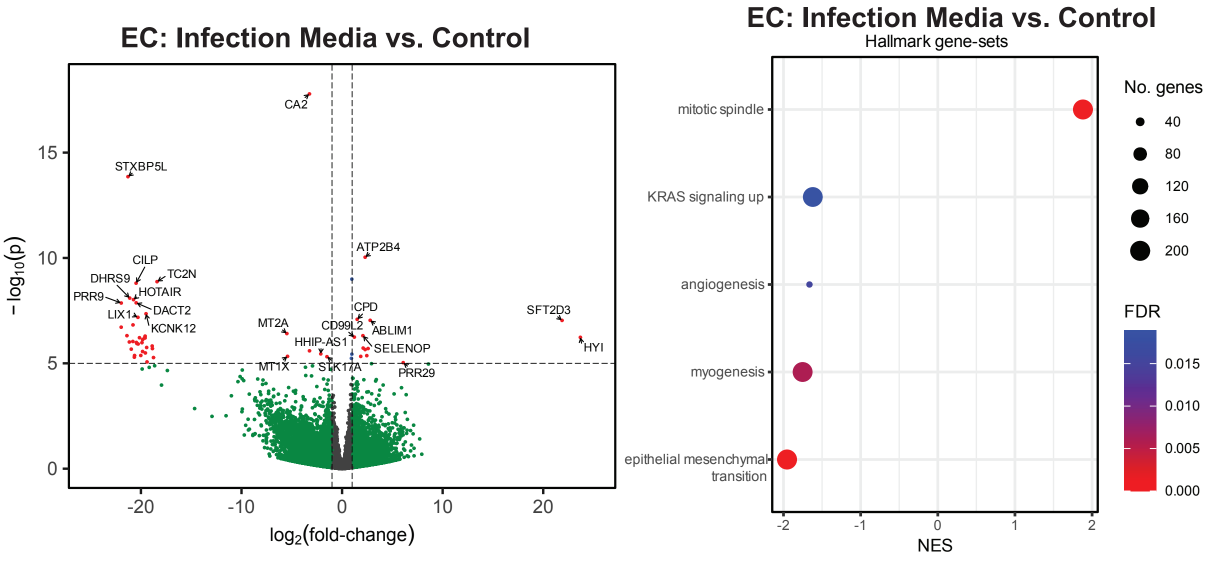


**Supplemental Fig 8: Exposure of ECs to SMC infection media does not induce inflammatory signaling.** ECs were treated with the media used for SMCs infections (Infection media) for 48 hours or maintained in their standard media. RNA was isolated and sequenced by bulk RNA sequencing. Volcano plots showing differential gene expression. Dot plots show gene-sets from the Hallmark collection[32] of the MSigDB that were enriched (FDR < 0.05) using gene-set enrichment analysis (GSEA)[76]. A full list of differentially expressed genes can be found in the Source Data file.

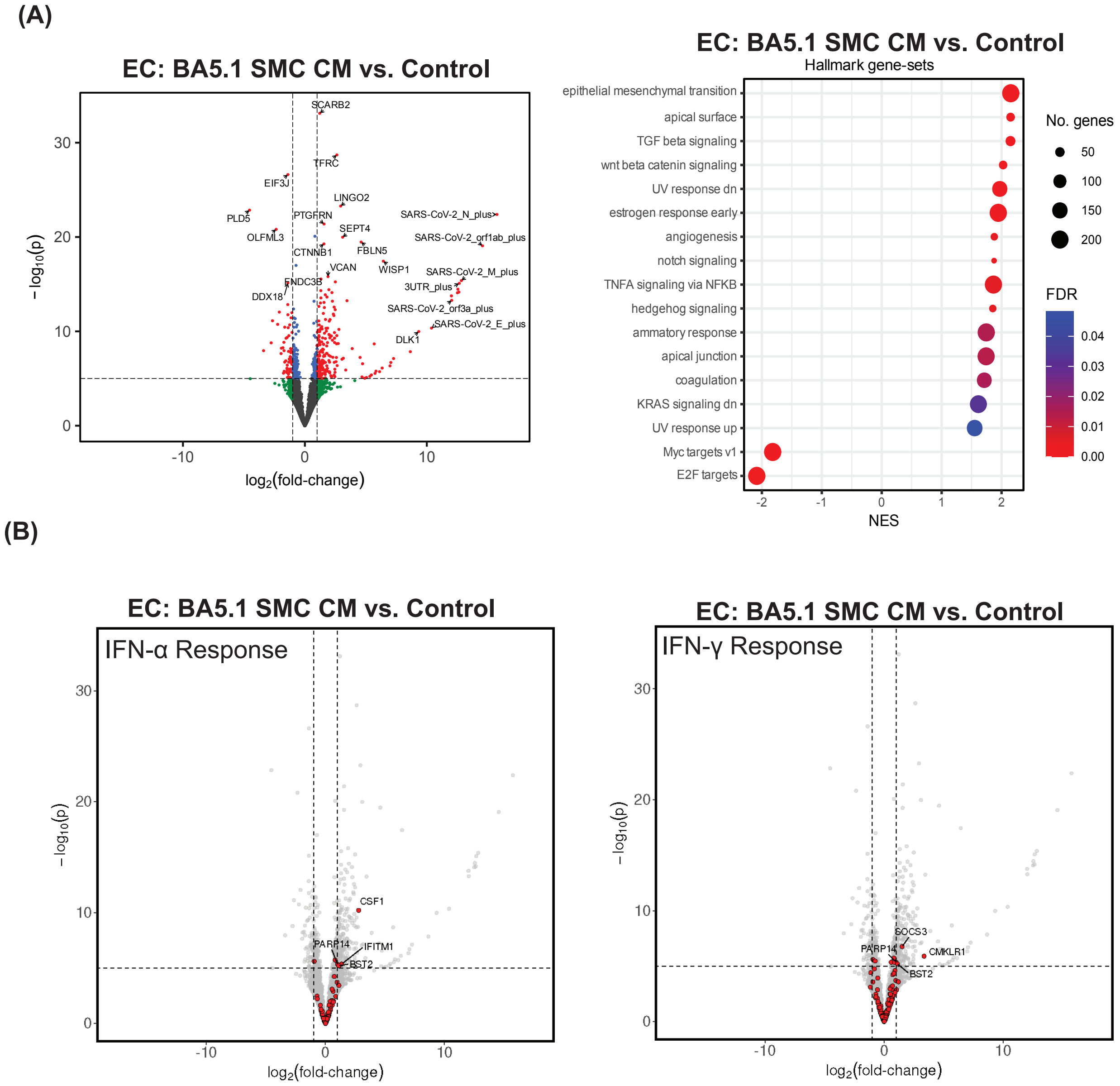


**Supplemental Fig 9: Paracrine signaling from BA5.1 infected SMCs does not induce significant inflammatory signaling in ECs.** Bulk RNA sequencing was performed on RNA isolated from ECs 48 hours after exposure to media from SMCs infected with the BA5.1 variant (MOI=1) (*BA5.1 SMC CM)* or control ECs *(Control)*. **(A)** (left) Volcano plots showing differential gene expression. (Right) Dot plots show gene-sets from the Hallmark collection[34] of the MSigDB that were enriched (FDR < 0.05) using gene-set enrichment analysis (GSEA). **(B)** Volcano plots with the genes from the “IFN-α response” (left) and

“IFN-γ response” (right) gene-sets of the MSigDB’s Hallmark collection[34] are highlighted in red, with GSEA “leading edge” genes indicated by name.

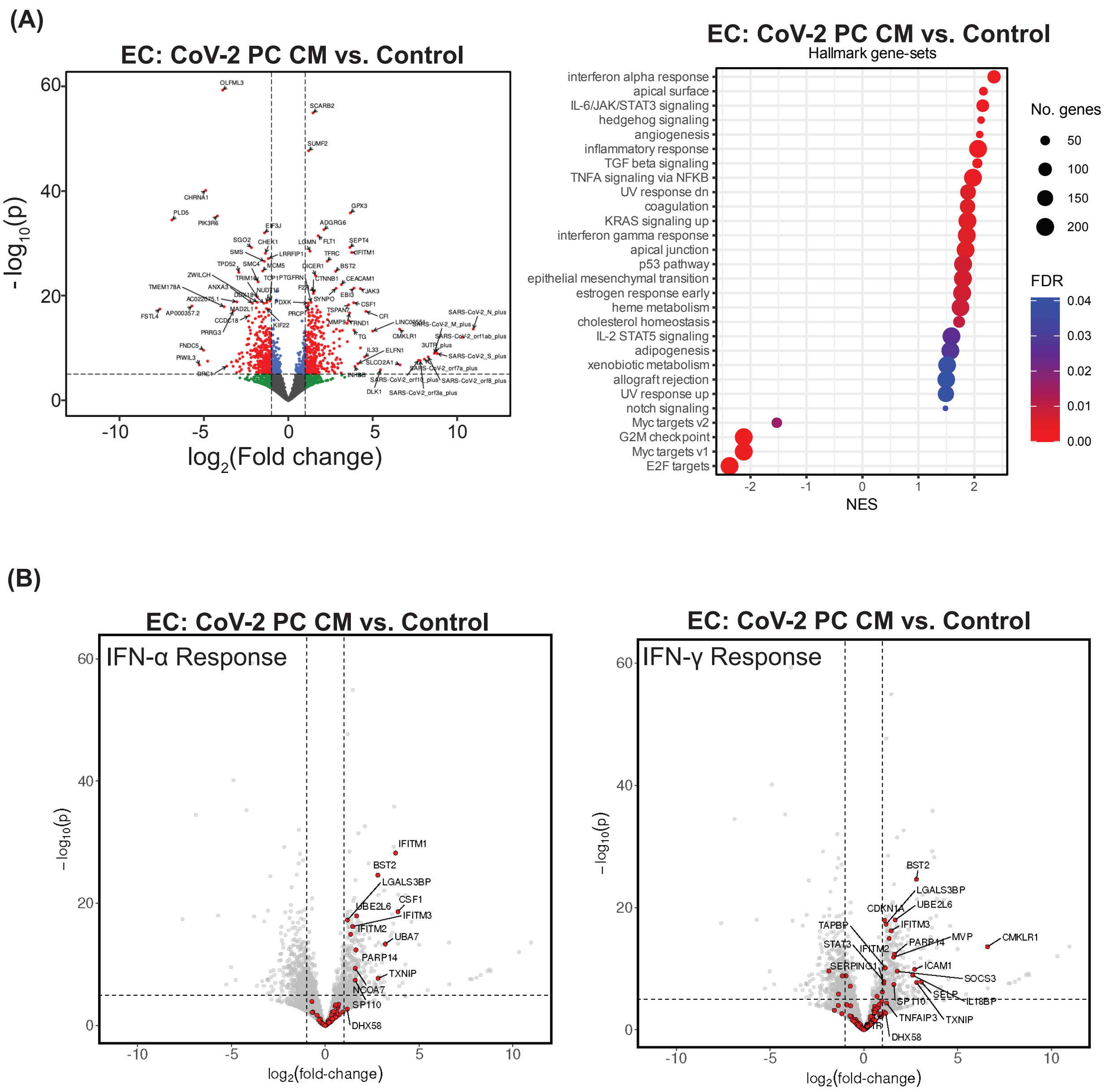


**Supplemental Fig 10: SARS-CoV-2 infected PCs release factors that promote inflammatory signaling in ECs.** Bulk RNA sequencing was performed on RNA isolated from ECs 48 hours after exposure PC-conditioned media. **(A)** ECs exposed to media from SARS-CoV-2 infected SMCs *(CoV-2 PC CM)* compared to control ECs *(Control)*. Volcano plots showing differential gene expression. Dot plots show gene-sets from the Hallmark collection[34] of the MSigDB that were enriched (FDR < 0.05) using gene-set enrichment analysis (GSEA) (**B)** Genes from the “IFN-α response” (left) and “IFN-γ response” (right) gene-sets of the MSigDB’s Hallmark collection[34] are highlighted in red, with GSEA “leading edge” genes indicated by name.

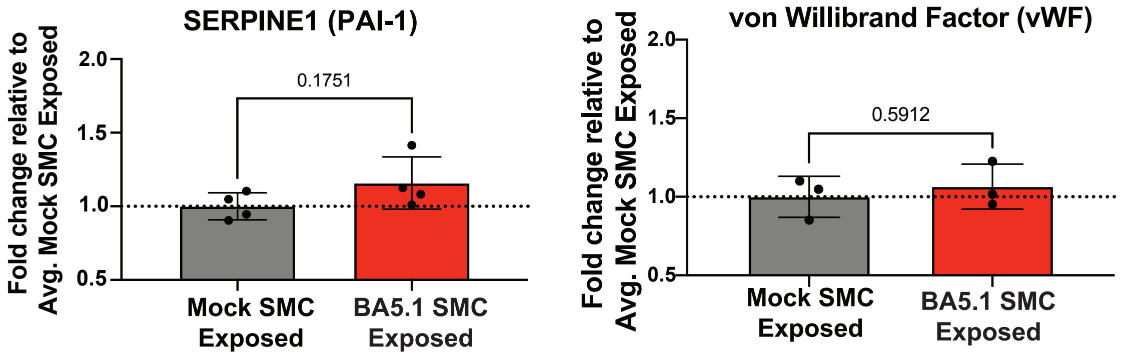


**Supplemental Fig. 11: Exposure of ECs to media from BA5.1 infected SMCs does not promote release of vWF or SERPINE1.** Quantitation of SERPINE1(PAI-1) and von Willebrand factor (vWF) in the media of ECs exposed to media from BA5.1 infected SMCs (*BA5.1 SMC Exposed*) for 48 hours or ECs exposed to media from mock infected SMCs *(Mock SMC Exposed)* for 48 hours. Four independent experiments were analyzed for SERPINE1 quantitation. Three independent experiments were analyzed for vWF quantitation. All values were plotted as fold change relative to the average value for *Mock SMC Exposed* samples). Bar graph shows mean value and error bar shows +/- SD. Conditions were compared using an unpaired t-test.

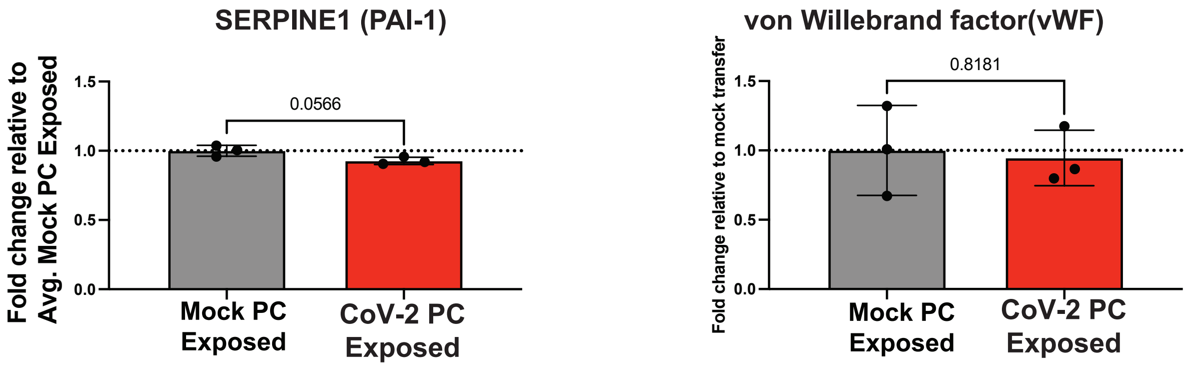


**Supplemental. Fig. 12: Exposure of ECs to media from SARS-CoV-2 infected PCs does not promote the release of vWF or SERPINE1.** Quantitation of SERPINE1(PAI-1) and von Willebrand factor (vWF) in the media of ECs exposed to media from SARS-CoV-2 infected PCs (*CoV-2 PC Exposed*) for 48 hours or exposed to media from mock infected PCs *(Mock PC Exposed)* for 48 hours. Three independent experiments were analyzed and results are expressed as a fold change relative to the average value recorded from *Mock PC Exposed* samples). Bar graph shows mean value and error bar shows +/- SD. Conditions were compared using an unpaired t-test.

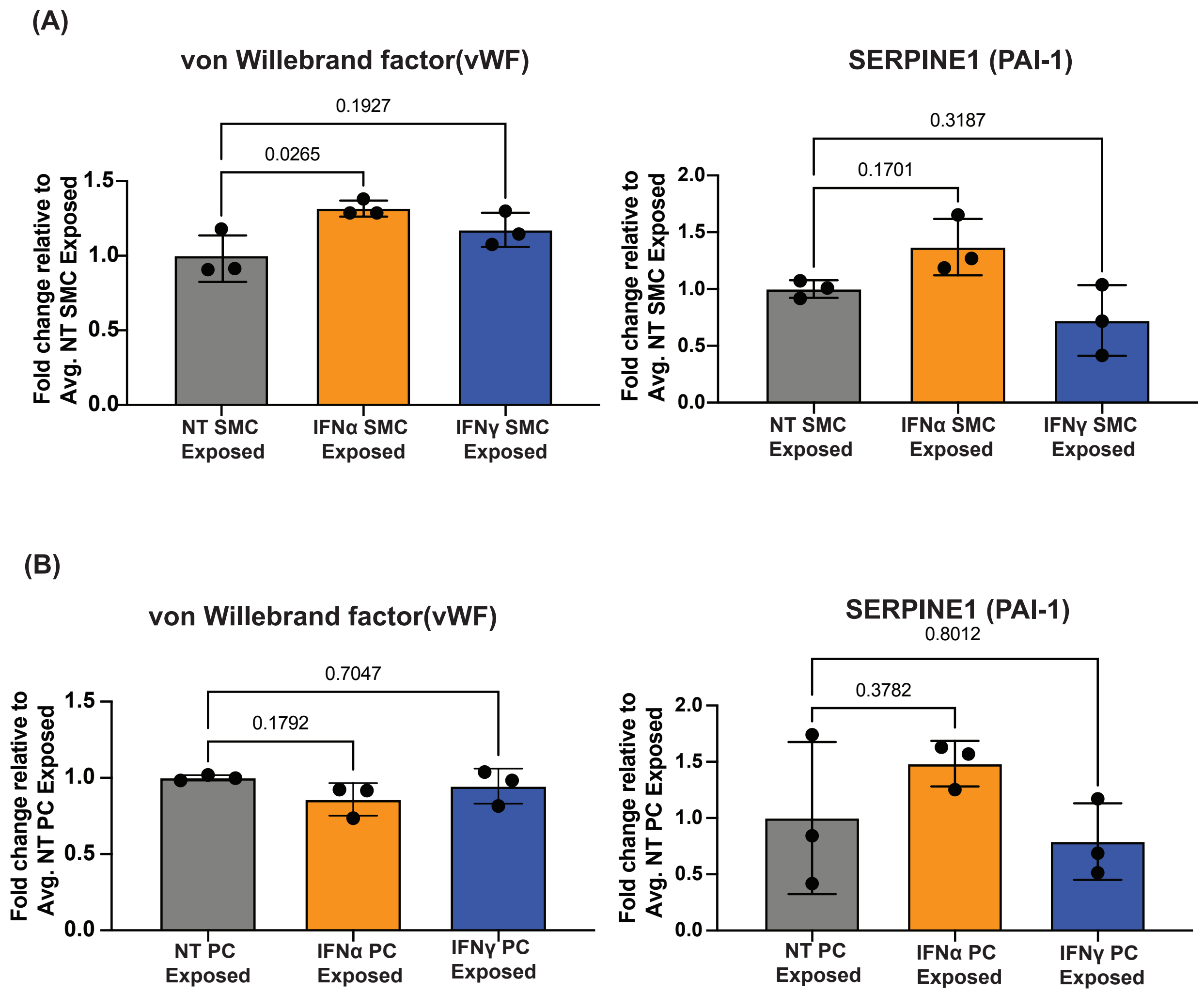


**Supplemental Fig. 13: Activation of inflammatory signaling in SMCs results in the release of factors that promote clotting cascades in ECs. (A)** hPSC derived SMCs were treated with 100U/ml IFNα or 20ng/ml IFNγ for 24 hours, cells were then washed twice and fresh media without IFN-α or IFN-γ was added. Media was then collected after 24 hours and added to hPSC derived ECs. Levels of vWF or SERPINE1 in the media were measured 48 hours later. Values from three independent experiments were used for vWF quantitation. Values from five independent experiments were used for SERPINE1 quantitation. All values are reported as a fold change relative to the average value for ECs treated with media from untreated SMCs (NT SMC Exposed). Bar graph shows mean value and error bar shows +/- SD. Conditions were compared using a one-way

ANOVA with Dunnett's multiple comparisons test, with a single pooled variance**. (B)** hPSC derived PCs were treated with 100U/ml IFNα or 20ng/ml IFN-γ for 24 hours, cells were then washed twice and fresh media without IFN-α or IFNγ was added. Media was then collected after 24hours and added to hPSC derived ECs. Levels of SERPINE1 or vWF in the media were measured 48 hours later. Three independent experiments were analyzed for vWF quantitation. Values from five independent experiments were used for SERPINE1 quantitation. All values are reported as a fold change relative to the average value for ECs treated with media from untreated PCs (NT PC Exposed). Bar graph shows mean value and error bar shows +/- SD. Conditions were compared using a one-way ANOVA with Dunnett's multiple comparisons test, with a single pooled variance**.**

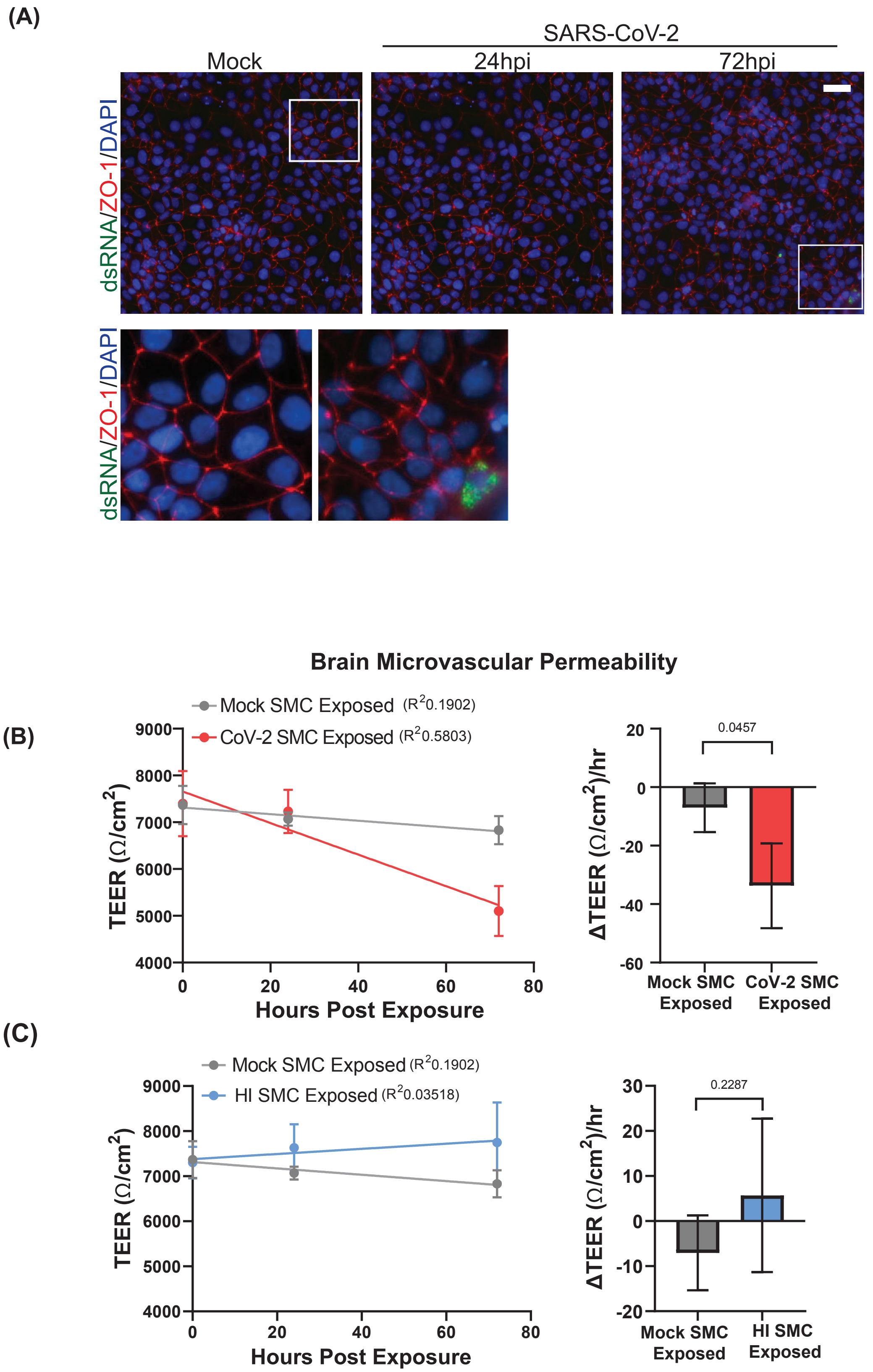
(A)
SARS-CoV-2
Mock
24hpi
72hpi
dsRNA/ZO-1/DAPI
dsRNA/ZO-1/DAPI
Brain Microvascular Permeability
(B)
Mock SMC Exposed (R2 0.1902)
CoV-2 SMC Exposed (R2 0.5803)
TEER (Ω/cm2)
Hours Post Exposure
0.0457
ΔTEER (Ω/cm2)/hr
Mock SMC Exposed
CoV-2 SMC Exposed
(C)
Mock SMC Exposed (R2 0.1902)
HI SMC Exposed (R2 0.03518)
TEER (Ω/cm2)
Hours Post Exposure
0.2287
ΔTEER (Ω/cm2)/hr
Mock SMC Exposed
HI SMC Exposed

**Supplemental Fig. 14: Exposure of hPSC-derived brain microvascular cells to SARS-CoV-2 or to media conditioned by SARS-CoV-2 infected SMCs.** **(A)** hPSC-derived brain microvascular endothelial cells (hBMECs) were exposed to live SARS-CoV-2 (MOI=0.1) for 24hr or 72hr hours. Cells were fixed and stained for the endothelial cell marker ZO-1 and dsRNA to detect viral replication. Lower images are enlarged images of regions highlighted by white boxes. The experiment was performed twice with similar results. Representative images from a single experiment are shown. **(B)** hBMECs were plated in trans-well plates and exposed to media from SMCs treated with heat-inactivated SARS-CoV-2 (HI) or media from uninfected SMCs (*Control)*. Trans-endothelial cell electrical resistance (TEER) was measured at 0, 24, and 72 hours after exposure. **(C)** Trans-endothelial cell electrical resistance (TEER) was measured at 0, 24, and 72 hours after exposure of hPSC-derived brain microvascular endothelial cells to media from SARS-CoV-2 infected SMCs (*Exposed*) or media from uninfected SMCs (*Control)*. For all experiments TEER was measure in three wells for each condition at each time point and the mean with standard deviation plotted.

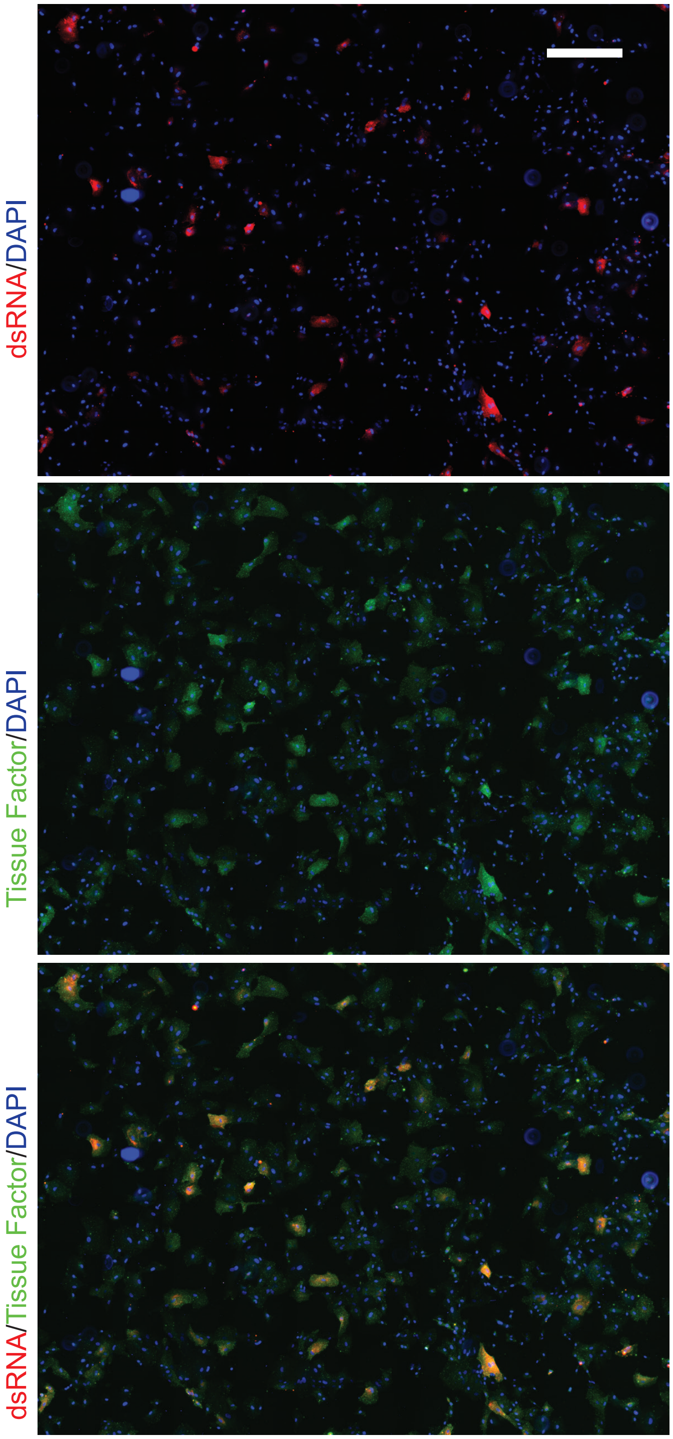


**Supplemental Fig. 15: Tissue factor staining is increased in cells with actively replicating SARS-CoV-2.** Representative immunofluorescence micrograph showing association of TF expression in infected SMCs at 48 hours post infection (MOI=0.1). The fraction of infected cells was determined by quantitating the number of dsRNA-positive cells and determined to be15.78% (155/982 DAPI+ cells). Scale bar = 200 μm

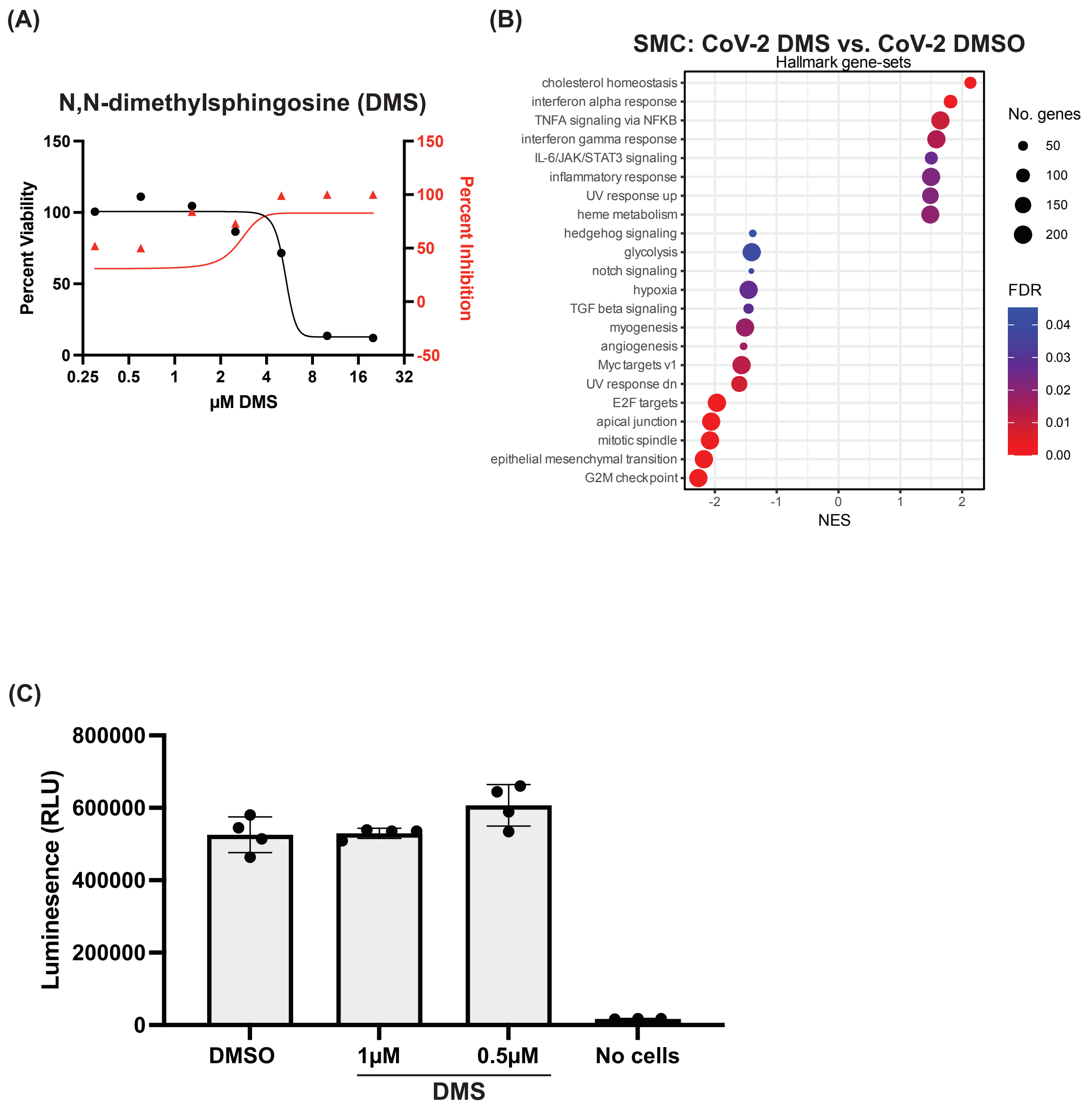

(A)
N,N-dimethylsphingosine (DMS)
Percent Viability
Percent Inhibition
µM DMS
0.25
0.5
1
2
4
8
16
32
150
100
50
0
-50
(B)
SMC: CoV-2 DMS vs. CoV-2 DMSO
Hallmark gene-sets
cholesterol homeostasis
interferon alpha response
TNFA signaling via NFKB
interferon gamma response
IL-6/JAK/STAT3 signaling
inflammatory response
UV response up
heme metabolism
hedgehog signaling
glycolysis
notch signaling
hypoxia
TGF beta signaling
myogenesis
angiogenesis
Myc targets v1
UV response dn
E2F targets
apical junction
mitotic spindle
epithelial mesenchymal transition
G2M checkpoint
NES
-2
-1
0
1
2
No. genes
50
100
150
200
FDR
0.04
0.03
0.02
0.01
0.00
(C)
Luminesence (RLU)
800000
600000
400000
200000
0
DMSO
1µM
0.5µM
No cells
DMS

**Supplemental Fig. 16: The SPHK inhibitor N,N-dimethyl-sphingosine reduces SARS-CoV-2 replication in hPSC-derived SMCs. (A)**SMCs infected with SARS-CoV-2 (MOI=0.1) were exposed to increasing amounts of N,N-dimethyl-sphingosine (DMS) during infection. The amount of infectious virus released into the media at 48 hours post-infection was quantitated by plaque assay. Cell viability was measured in uninfected SMCs at the corresponding time point and DMS dose. **(B)** Dot plot summary of GSEA comparing SARS-CoV-2 infected SMCs with and without exposure to DMS (1uM) during infect. The Hallmark collection[34] of gene-sets from the MSigDB was used and gene-sets were plotted only when FDR < 0.05 for enrichment in at least one of the between-condition comparisons. **(C)** Viability of ECs following 48 hours of exposure to 1uM or 0.5uM DMS was quantitated by CellTiter Glo assay. Bar graph shows mean value and error bar shows +/- SD.

Supplemental Table 1: Summary of antibodies used in this study

| **Primary Antibodies** | **Source** | **Catalog Number** | **Dilution** |
|---|---|---|---|
| VE-Cadherin | R&D Systems | AF938 | 1:250 |
| vWF | Abcam | ab6994 | 1:250 |
| PECAM1(CD31) | Abcam | ab9498 | 1:250 |
| SMA | Abcam | ab5694 | 1:200 |
| PDGFR$\beta$ | Cell Signaling | 3169S | 1:200 |
| NG2 | Invitrogen | 14-6504-82 | 1:200 |
| dsRNA (J2) | Novus | NBP3-11395 | 1:2000 |
| Tissue Factor | Abcam | ab228968 | 1:250 |
| ZO-1 | Life Technologies | 402200 | 1:200 |
| VE-Cadherin-PE (Flow for EC) | Invitrogen | 12-1449-82 | 1:50 |
| PECAM1-AlexaFluor 647 (Flow for EC) | Abcam | Ab215912 | 1:50 |
| SMA-AlexaFluor 594 (Flow for SMC) | Cell Signaling | 36110S | 1:50 |
| PDGFRb-APC (Flow for SMC) | Abcam | Ab119861 | 1:50 |
| NG2-APC (Flow for PC) | R&D Systems | FAB2585A | 1:25 |
| PDGFRb-APC (Flow for PC) | BioLegend | 323512 | 1:25 |
| PDGFRa-PE (Flow for PC) | BioLegend | 323606 | 1:25 |
| Secondary Antibodies | Source | Catalog Number | Dilution |
| Mouse-488 | Life Technologies | A21202 | 1:1000 |
| Mouse-568 | Life Technologies | A10037 | 1:1000 |
| Mouse-647 | Life Technologies | A31571 | 1:1000 |
| Rabbit-488 | Life Technologies | A21206 | 1:1000 |
| Rabbit-568 | Life Technologies | A11011 | 1:1000 |

| | | | |
|---|---|---|---|
| Rabbit-647 | Life Technologies | A31573 | 1:1000 |
| Goat-488 | Life Technologies | A11055 | 1:1000 |
| Goat-568 | Life Technologies | A11057 | 1:1000 |
| Goat-647 | Life Technologies | A21447 | 1:1000 |

Supplemental Table 2: Media formulations for differentiation and infection media

| | Final Concentration | Source | Catalog # |
|---|---|---|---|
| **MelM** | | | |
| E6 | | Thermo Fisher Scientific | A1516401 |
| L-Ascorbic acid 2-phosphate sesquimagnesium salt hydrate (AA) | 60ug/ml | Sigma | A8960-5G |
| CHIR 99021 | 8uM | Biogems | 2520691 |
| BMP4 | 25ng/ml | Peprotech | 120-05ET |
| **EC1** | | | |
| E6 | | Thermo Fisher Scientific | A1516401 |
| Forskolin | 2nM | Biogems | 6652995 |
| AA | 60ug/ml | Sigma | A8960-5G |
| VEGF | 200ng/ml | Peprotech | 100-20-50µg |
| CP-673451 | 2nM | Selleck | S1536 |
| SB 431542 | 10uM | Biogems | 3014193 |
| **EC2** | | | |
| hESFM | | Thermo Fisher Scientific | 11111044 |
| Forskolin | 2nM | Biogems | 6652995 |
| AA | 60ug/ml | Sigma | A8960-5G |
| VEGF | 200ng/ml | Peprotech | 100-20-50µg |
| CP-673451 | 2nM | Selleck | S1536 |
| SB 431542 | 10uM | Biogems | 3014193 |
| **EC3** | | | |
| hESFM | | Thermo Fisher Scientific | 11111044 |
| AA | 60ug/ml | Sigma | A8960-5G |
| VEGF | 200ng/ml | Peprotech | 100-20-50µg |
| CP-673451 | 2nM | Selleck | S1536 |
| SB 431542 | 10uM | Biogems | 3014193 |
| Exherin (ADH-1) | 25ug/ml | AdooQ BioScience | A13689 |
| **EC4** | | | |
| hESFM | | Thermo Fisher Scientific | 11111044 |
| B27 | 1:50 | Thermo Fisher Scientific | 17504044 |
| EGF | 20ng/ml | Peprotech | AF-100-15 |
| Heparin | 2ug/ml | StemCell Technologies | 07980 |
| VEGF | 20ng/ml | Peprotech | 100-20-50µg |
| CP-673451 | 2nM | Selleck | S1536 |
| SB 431542 | 10uM | Biogems | 3014193 |
| S1P | 5nM | Sigma | S9666-1MG |
| **EC5** | | | |
| hESFM | | Thermo Fisher Scientific | 11111044 |
| B27 | 1:50 | Thermo Fisher Scientific | 17504044 |
| EGF | 20ng/ml | Peprotech | AF-100-15 |
| Heparin | 2ug/ml | StemCell Technologies | 07980 |

| | | | |
|---|---|---|---|
| VEGF | 20ng/ml | Peprotech | 100-20-50µg |
| bFGF | 50ng/ml | Peprotech | 100-18B |
| **PC1/SMC1** | | | |
| E6 | | Thermo Fisher Scientific | A1516401 |
| Forskolin | 2nM | Biogems | 6652995 |
| AA | 60ug/ml | Sigma | A8960-5G |
| VEGF | 200ng/ml | Peprotech | 100-20-50µg |
| **PC2/SMC2** | | | |
| hESFM | | Thermo Fisher Scientific | 11111044 |
| Forskolin | 2nM | Biogems | 6652995 |
| AA | 60ug/ml | Sigma | A8960-5G |
| VEGF | 200ng/ml | Peprotech | 100-20-50µg |
| **PC3** | | | |
| hESFM | | Thermo Fisher Scientific | 11111044 |
| AA | 60ug/ml | Sigma | A8960-5G |
| VEGF | 200ng/ml | Peprotech | 100-20-50µg |
| SB 431542 | 10uM | Biogems | 3014193 |
| **PC4** | | | |
| hESFM | | Thermo Fisher Scientific | 11111044 |
| B27 | 1:50 | Thermo Fisher Scientific | 17504044 |
| EGF | 20ng/ml | Peprotech | AF-100-15 |
| Heparin | 2ug/ml | StemCell Technologies | 07980 |
| SB 431542 | 10uM | Biogems | 3014193 |
| PDGFbb | 10ng/ml | Peprotech | 100-14B-10UG |
| **SMC3** | | | |
| hESFM | | Thermo Fisher Scientific | 11111044 |
| AA | 60ug/ml | Sigma | A8960-5G |
| VEGF | 200ng/ml | Peprotech | 100-20-50µg |
| CP-673451 | 2nM | Selleck | S1536 |
| **SMC4** | | | |
| hESFM | | Thermo Fisher Scientific | 11111044 |
| B27 | 1:50 | Thermo Fisher Scientific | 17504044 |
| EGF | 20ng/ml | Peprotech | AF-100-15 |
| Heparin | 2ug/ml | StemCell Technologies | 07980 |
| CP-673451 | 2nM | Selleck | S1536 |
| **SMC5** | | | |
| hESFM | | Thermo Fisher Scientific | 11111044 |
| B27 | 1:50 | Thermo Fisher Scientific | 17504044 |
| EGF | 20ng/ml | Peprotech | AF-100-15 |
| Heparin | 2ug/ml | StemCell Technologies | 07980 |
| TGFβ | 10ng/mlM | Peprotech | 100-21-500ug |
| **Infection Media** | | | |

| | | | |
|---|---|---|---|
| hESFM | | Thermo Fisher Scientific | 11111044 |
| B27 | 1:50 | Thermo Fisher Scientific | 17504044 |
| EGF | 20ng/ml | Peprotech | AF-100-15 |
| Heparin | 2ug/ml | StemCell Technologies | 07980 |
| VEGF | 20ng/ml | Peprotech | 100-20-50µg |
| bFGF | 50ng/ml | Peprotech | 100-18B |
